# Sedimentary ancient DNA analysis from the Iron Age courtyard and cooking pit site of Rødskjær in Troms, Northern Norway

**DOI:** 10.64898/2026.10.30.763041

**Authors:** Mary Lucas, Mikael Cerbing, Claire-Elise Fischer, Stephen Wickler, Inger Greve Alsos, Youri Lammers, Galina Gusarova, Marieke Beaulieu, Antony G Brown

## Abstract

Courtyard sites, which are Iron Age structures unique to Norway, are often associated with cooking pit fields suggesting feasting practices, but little is known about the main purpose of these sites. To address this, a framework for sedimentary ancient DNA (sedaDNA) metabarcoding sampling was undertaken at the site of Rødskjær, northern Norway. Of the 63 sediment samples from cooking pits, fire pits, a hearth, a fireplace, the natural shell-sand and topsoil, 52 successfully detected plant and animal sedaDNA. The plant diversity was high with 41-100 taxa. SedaDNA identified archaeologically significant taxa that were not detected by other methods, including *Avena* (oats), *Quercus* (oak), *Equus* (horse), and *Salmo salar* (salmon). Additionally, from inter-sample comparison, and with reference to worm sedaDNA we identified the archaeological contexts that were affected by partial bioturbation and root penetration that restricted interpretation of spatial or temporal differences in use of these features. This pioneering study shows the importance of taking many samples from modern and natural sediments as well as using multi-proxy analysis when conducting sedaDNA metabarcoding studies at open-air archaeological sites. If these factors are taken into consideration, sedaDNA from cooking pits can be a valuable resource for investigating uses of courtyard and cooking pit field sites.

## Introduction

Courtyard sites have been found along the Norwegian coast from Agder to Bjarkøy in northern Norway (Figure 1). The earliest courtyard sites were constructed in the early Roman Iron Age (1-200 AD) in southwestern Norway and spread to northern Norway in the late Roman Iron Age (200-400 AD) (Grimm, 2010). They usually include a cluster of houses often built in a circle or a semicircle around an open central space. The number of houses at these sites range from four to eighteen and the houses are of similar size and construction. The sites were commonly used for hundreds of years. Some might still have been active during the early Viking Age (800-1050 AD), but most appear to have fallen out of use during the early Merovingian Period (550-750 AD) (Olsen, 2005; Storli, 2006; Grimm, 2010; Hatling and Olsen, 2012; Cerbing, 2025). While these sites are important to the understanding of socio-political structures in Norway during much of the Iron Age, surprisingly little is known about their associated activities and economies.

**Figure 1:**
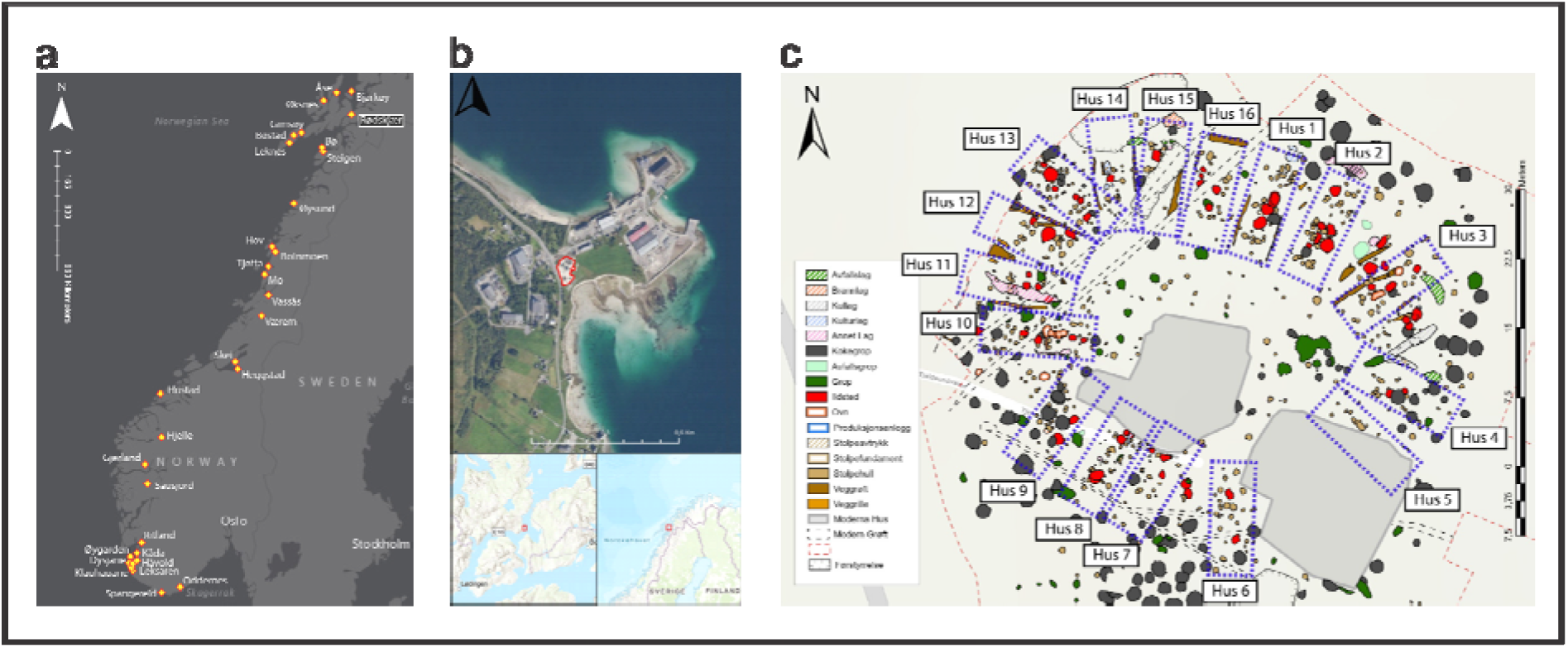
**a)** Location of Rødskjær in relation to all documented courtyard sites found in Norway. Location of Rødskjær highlighted in black. **b)** Rødskjær location and excavation area outlined in red. **c)** Site drawing with identified house structures from the courtyard period of the site.

Clusters of cooking pit features, referred to here as “fields”, are frequently associated with courtyard sites (Johansen and Søbstad, 1978; Martens, 2005; Hatling and Olsen, 2012; Cerbing, 2025). The term cooking pit is used to describe features that have been dug out to build a fire and filled with heated rocks to produce steam or to warm water and assumed to be used for cooking (Gundersen et al., 2020). Despite the frequent presence of cooking pit fields at courtyard sites, their precise relationship to these sites remains poorly understood. The use of cooking pits for preparation of large quantities of food in association with feasting and other activities at the courtyard sites has been proposed by a number of researchers (Gjerpe, 2001; Narmo, 2009; Ødegaard, 2021). Radiocarbon dates and stratigraphy from the two most recent courtyard site excavations suggest the cooking pit fields appeared earlier than the courtyard site itself (Hatling and Olsen, 2012; Cerbing, 2025). It is possible that the same is true for other courtyard sites, but it is difficult to assess this from the primary documentation (Sjøvold 1971; Johansen and Søbstad, 1978).

Sedimentary ancient DNA (sedaDNA) has the potential to provide information about the use history and purpose of these sites and whilst it has been used extensively for lake cores and cave sites, open-air archaeological sites are just beginning to be explored for sedaDNA investigation (Brown et al., 2025; Han et al., 2025; La Torre et al., 2026). This is partly due to the different methodological challenges they pose compared to protected sites such as lakes and caves (La Torre et al., 2026). However, studies investigating sedaDNA deposition and mineralogical interaction are addressing these challenges (Hebsgaard et al., 2009; Seersholm et al., 2016; Gu et al., 2023; Zampirolo et al., 2024; Sawafuji et al., 2025; La Torre et al., 2026). Additionally, there has also been little research on heated archaeological features due to the assumption that sedaDNA would not preserve through the high temperatures of the heating process (Ottoni et al., 2009; Braadbaart et al., 2020). However, uncharred food waste is often found in these archaeological features suggesting incomplete combustion and relatively low temperatures. There is also the potential for sedaDNA to preserve after the burning event of the feature through backfill secondary deposits (Wygel et al., 2024). For this reason, the sediments within the cooking pits could also be suitable environments for sedaDNA preservation (Özdoğan et al., 2024).

This study uses sedimentary ancient DNA metabarcoding of animal and plant DNA from heated archaeological features to investigate dietary and site use practices at Rødskjær. Metabarcoding is a targeted sedaDNA sequencing approach with high success of identifying taxa to the species level compared to other methods and therefore has a higher chance of identifying species composition and species turnover over time at the site (Strandberg et al., 2026). Cooking pits, as well as fire pits, a fireplace, and a hearth were sampled. Fire pits, fireplaces, and hearths are identified separately from cooking pits as they are built on the ground surface or prepared substrate whereas fire pits are constructed below the ground surface (Mentzer, 2017). Sampling these features allows for a comparison of heated features at the site that might have been used for different purposes and burnt at different temperatures potentially with different fuel sources. The results of sedaDNA analyses are compared with macrofossil, charcoal and zooarchaeological analyses to understand what new information sedaDNA can contribute to our understanding of these sites.

## 2. Materials and methods

### 2.1 Rødskjær excavation and site background

The courtyard site at Rødskjær, on the eastern side of the island Hinnøya in northern Norway 230 km north of the arctic circle (Figure 1), was excavated in 2023 and 2024 by the Arctic University Museum of Norway due to planned commercial development of the area (Cerbing, 2025). The site, which is situated close to the shoreline along the Tjeldsund Strait, represents a landscape utilized by people throughout prehistory (Hole and Mikalsen, 2016; Melsæther, 2016; Bruun and Oppvang, 2025). The site was constructed on a small coastal promontory (spit or ness) c. 430 m long and 210 m wide, with a maximum elevation of 7 m above sea level. At the time of excavation, the site was a residential area with two houses, a shed, a meadow, and a modern garden (Figure S1 and S2). Based on the design the shed was most likely first functioned as a barn.

Archaeological intervention methods involved mechanical surface stripping followed by manual excavation of intact cultural deposits. Radiocarbon dating, phosphate analysis, zooarchaeological, and macrofossil analyses were carried out after the excavation (Cerbing, 2025). Radiocarbon dates suggest the use of the cooking pits began in the last century BC with construction of the courtyard site in the early third century AD. Additionally, many of the cooking pits stratigraphically predate construction of the courtyard site. In total, 155 cooking pits were documented at the site in an area of about 3400 m^2^. During excavation additional cooking pits were identified outside the main site area.

The highest level of site activity appears to be during the period 200-400 AD with continued intensive use up until 600 AD, although there are fewer dates from the period 400-600 AD. This was followed by more sporadic activity up until the early Viking Age (800-900 AD) including finds of two Viking brooches (Cerbing, 2025). Whether the lower number of dates from the period 400-600 AD is due to decreased site activity or modern destruction of later cultural layers is difficult to determine, however, it is clear that the courtyard site was established in the third century AD and went into disuse by the end of the 6th/early 7th century AD.

At the peak of activity, the Rødskjær site might have had up to 16 houses, and almost all houses had signs of modification including repairs and reconstruction (Cerbing, 2025). The houses were similar in size and appearance, as is common in courtyard sites (Johansen & Søbstad, 1978; Grimm, 2010; Hatling & Olsen, 2012) and appear to be roughly 12 m long and 4 m wide and were placed in an oval around the central courtyard, probably with the entrance towards the courtyard. The houses were aisled with twin wooden posts holding up the roof, and they probably had wattled walls and some may have had earthen walls between the houses, but the site had been plowed out, removing any possible remains of earthen walls and no definite proof of this could be found.

Except for faunal remains, very few finds were documented at the site. Most of the artifacts were discovered in the surface stripped topsoil layer. The artifacts were mostly small personal items such as brooches, needles, and whetstones. Of the faunal remains identified, 89% are mammals, and 90% of these are *Ovis* (sheep) and/or *Capra* (goat) (Cerbing, 2025). The low quantity of fish bones found in the zooarchaeological assemblage is unusual for coastal residential sites in northern Norway during this period (Naumann et al., 2014; Niemi, 2022). Almost none of the bones had signs of burning, suggesting that most, if not all, food preparation took place in the cooking pits. This indicates that the eating practices at Rødskjær were focused on certain types of food, prepared in a specific way, unlike the food practices commonly associated with residential sites (Bukkemoen, 2017).

### 2.2 Field Sampling

SedaDNA samples were collected from features across the courtyard site and cooking pit field to account for any spatial variability in DNA deposition. Samples were also taken from the topsoil and natural sediments surrounding the features to identify differences in species composition between archaeological and naturally occurring sediments and soils. In total, 63 samples were taken from nine features at the site (Figure S3-S11). The two natural shell-sand sediments were sampled adjacent to features 9720 and 11450. Because the site was surface stripped before sedaDNA samples could be taken, the two topsoil samples were taken adjacent to the site area (Figure 2). All precautions were taken to avoid modern and cross-contamination. Samples were collected with personal protective equipment (gloves, mask, hairnet) and sampling equipment was cleaned with bleach and ethanol between each sample. The outer layer of sediment was removed three times with a new, cleaned knife and then samples were collected in sterile 15ml falcon tubes pressed into the sediment profile and stored at +2-5 ℃ before laboratory analysis.

**Figure 2:**
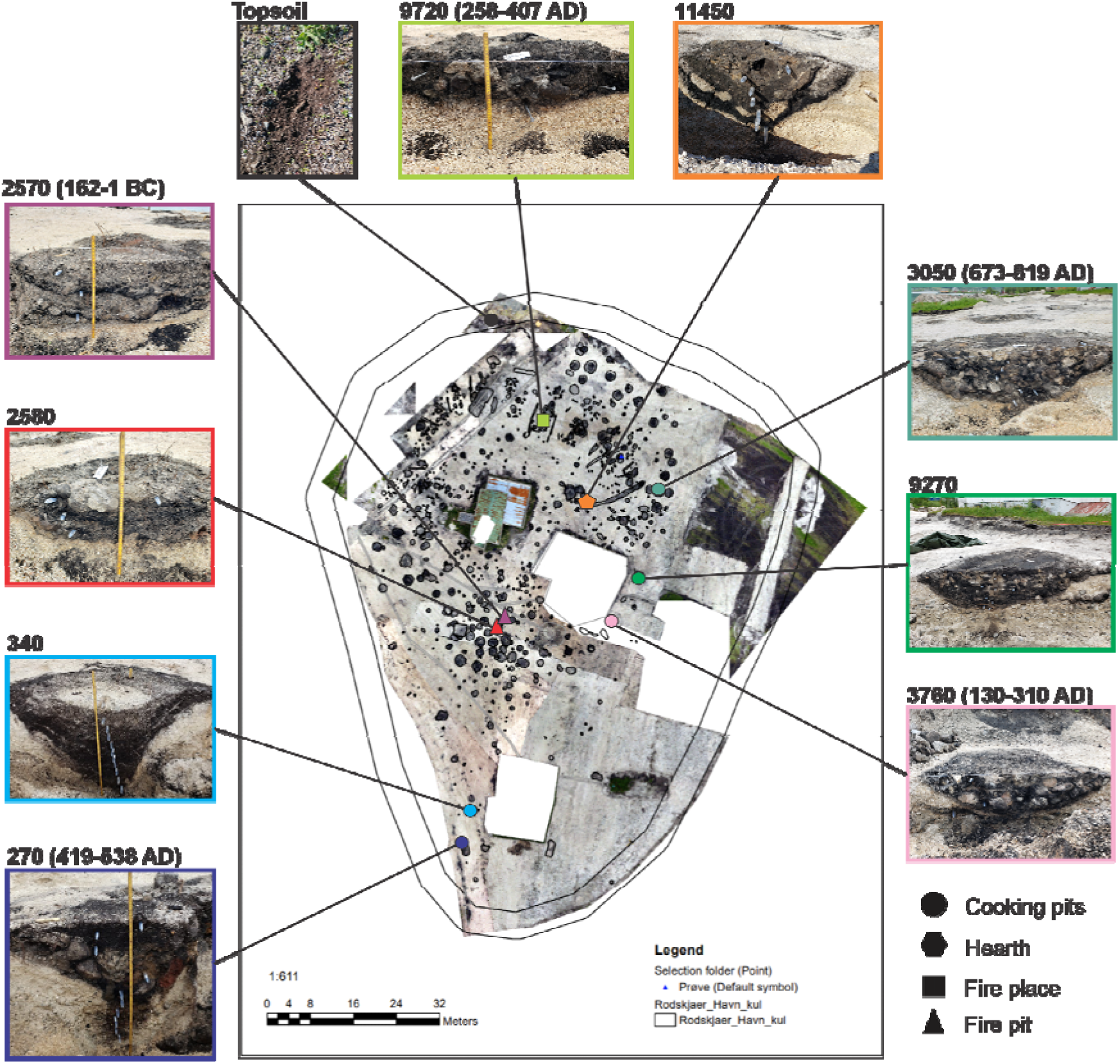
Overview of archaeological excavation at Rødskjær highlighting the archaeological features sampled for this study. Radiocarbon dates were taken from five of the features and are displayed next to the feature name. Shapes represent feature type and color identifies the individual features in figure 3 and supplementary figures.

### 2.3 Ancient DNA analysis

Sediment samples were extracted in a dedicated ancient DNA laboratory at The Arctic University Museum of Norway. Between 0.25-0.35g of sediment was subsampled for sedaDNA extraction alongside a negative control sample using a modified Qiagen DNeasy PowerSoil PowerLyzer (Qiagen Norge, Oslo, Norway) protocol (Voldstad et al., 2020). Four negative controls were used and each sample and control was amplified in eight replicates in a PCR laboratory located in a separate building from the ancient DNA laboratory. To detect plant sedaDNA, samples were amplified with primers amplifying the *trn*L P6 loop region of the chloroplast genome (Taberlet et al., 2007). This region of the chloroplast is targeted because it is suitable for high resolution identification of very short fragments of sedaDNA (Taberlet et al., 2007). For the retrieval of animal sedaDNA, primers amplifying the mitochondrial 16S locus were used (Giguet-Covex et al. 2014). The mitochondrial 16S rRNA region is proven to identify short DNA fragments of mammalian DNA to a high resolution (Vences et al., 2005). Whilst the 16S primers are designed for mammalian DNA, they can also detect other taxa such as fish, birds, amphibians, and invertebrates in what is referred to as ‘bycatch’. A forward and reverse blocking primer was used to reduce the amplification of human sedaDNA (Garcés-Pastor et al. 2022). Both plants and mammals were amplified using uniquely dual tagged primer sets (Supplementary data 2 and 3). After amplification, the samples were pooled, cleaned and converted into eight DNA libraries with the Illumina TruSeq DNA PCR-Free protocol (Illumina Inc., CA, USA). The libraries were then sequenced at ∼10% of 2 × 150-cycle mid-output flow cell on the Illumina NextSeq platform at Genomics Support Centre Tromsø, UiT.

The sequenced DNA libraries were processed through a bioinformatic pipeline constructed using the OBITools software package (Boyer et al., 2016). The reference databases used to process the plant sequences were, in order of prioritized matches; PhyloNorway (Alsos et al., 2020, Wang et al. 2021), Arctborbryo (Sønstebø et al., 2010; Willerslev et al., 2014; Soininen et al., 2015), PhyloAlps (Garcés-Pastor et al., 2022), and EMBL release 143. EMBL release 143 was also used to process the mammal sequences. These databases provide reference DNA sequences that can then be used for taxonomic annotation of the obtained sedaDNA. Results were filtered to remove low-quality data and possible false positives. Plant identifications were kept if they had 100% match to one of the reference libraries used. Plant data was filtered with a minimum of one repeat and three reads per sequence in a sample and a minimum of three repeats and ten reads across the entire data set. We retained >95% match to reference data of mammal identifications with a minimum of one repeat and ten reads per sequence in a sample and a minimum of one repeat and 75 reads across the entire data set. DNA sequences that were found in the negative controls and were suspected of being a contaminant were removed (Supplementary data 7 and 11). Identifications were cross-checked and a final taxonomic assignment was made using reference databases, manual BLASTing, and species occurrence data. If the same taxon was identified by multiple DNA sequences, the sequences were collapsed together and considered as one result in further analysis. All species detections were plotted with the R package *Rioja* (version: 1.0-7) (Juggins, 2019) in R/RStudio version 4.6.1 (R Core Team 2026) (Figures S12-S23).

### 2.6 Statistical analysis

Patterns of plant species presence between features were compared through unconstrained ordination: Principal Component Analysis (PCA) performed using presence/absence data of the plant species and samples grouped by feature. Hellinger transformation (Legendre and Gallagher, 2001) was applied to the species presence/absence data matrix prior to the ordination. Both data transformation and the PCA analyses were carried out in *Vegan* package (Dixon, 2003) R/RStudio and the plotting was carried out using IBM SPSS Statistics software (Version: 29.0.0.0). To facilitate interpretation of the sample groupings we produced factor loading plots visualizing the relationships of the variables (plant taxa) with the Principal Components (PC1 and PC2).

PCoA and PERMANOVA (Permutational Multivariate Analysis of Variance) analyses were run to evaluate if the between feature variability was greater or less than the within feature variability. Analysis was performed with the *vegdist()* function in the R package *Vegan* using Jaccard similarity matrix (Real and Vargas, 1996). PERMANOVA analysis was performed using the adonis function with 999 permutations *Vegan*. The analyses were run first with all plant species and then again after removing the species found in the topsoil to assess temporal differences in deposition of species not found in the topsoil. Features with less than five samples were excluded from analysis.

We additionally explored whether the detection of deep burrowing (anecic and endogeic) worms was significantly associated with the presence of domesticate species, which could be indicative that species assemblages observed at depth are impacted by bioturbation of material from surface soil layers (Armour-Chelu and Andrews, 1994). As worm detection was a bycatch of the mammalian target primer (Lammers et al., 2019), we considered only presence-absence taxa data. We used the method developed by Veech (2013) using a combinatory approach to compare the observed co-occurrence of species pairs (i.e. number of sites where they are found together) with their expected co-occurrence, defined as the product of each species probability of occurrence multiplied by the number of sampling sites. The analysis was run using the R-cooccur package (Griffith et al., 2016), calculating co-occurrence probabilities using the hypergeometric distribution and excluding species pairs that were expected to have less than one co-occurrence.

## 3. Results overview

Of the 63 samples, sedaDNA was successfully extracted from 52 (82.5%) samples with 11 (17.5%) not yielding sedaDNA (Supplementary data 1). A total of 15,144,536 and 906,540 post filter reads were identified for plants and mammals respectively. Cooking pits and the hearth feature contained the highest number of taxa on average (Table 1). Species richness within the features was significantly higher than the natural shell-sand with 39 taxa per sample compared to 16 taxa per sample.

**Table 1.** Details and results from features sampled.

| Feature ID | Feature Type | Dimensions | Date (calBC/ calAD) | # of samples | # successful | # of Plant Taxa | # of Animal Taxa | Average # Taxa Per Sample |
|---|---|---|---|---|---|---|---|---|
| Topsoil | - | 0-5cm depth | Modern | 2 | 1 | 69 | 12 | 81 |
| Shell sand | - | - | - | 2 | 2 | 16 | 7 | 16 |
| 340 | Cooking pit | 185 dia x 150cm d | - | 11 | 4 | 100 | 15 | 60.8 |
| 9270 | Cooking pit | 228cm dia x 36cm d | - | 10 | 10 | 77 | 10 | 27.1 |
| 3050 | Cooking pit | 212cm dia x 48cm d | 673-819 AD | 8 | 8 | 96 | 20 | 40.6 |
| 270 | Cooking pit | 340 x 150cm –140cm d | 419-538 AD | 10 | 8 | 77 | 16 | 44 |
| 3760 | Cooking pit | 152cm dia x 41cm d | 130-310 AD | 3 | 3 | 66 | 14 | 47.7 |
| 11450 | Hearth | 74cm dia x 46cm d | - | 7 | 7 | 83 | 9 | 54.6 |
| 9720 | Fire place | 152cm dia x 19cm d | 258-407 AD | 3 | 2 | 41 | 6 | 34 |
| 2580 | Fire pit | 140 x 135cm –20cm d | - | 4 | 4 | 53 | 3 | 27.5 |
| 2570 | Fire pit | 150 x 120cm –36cm d | 162-1 BC | 3 | 3 | 41 | 6 | 18.3 |

### 3.1 Plant results

The plant species detected included 72 taxa of forbs, 17 graminoids, 13 trees and shrubs, six dwarf shrubs, nine vascular cryptograms (ferns, horsetails, clubmosses), five bryophytes, and one aquatic taxon. In total, we detected 123 plant taxa across the 52 samples (Figure S12-S19, supplementary data 7). Of these, 40 were identified to genus level and 72 (59%) to species level. There were on average 36 plant taxa present per sample. The site is comprised mostly of native vascular plants typical of coastal meadow and open forest habitats in the study area: forbs, trees/shrubs, and graminoids (grasses, sedges and rushes). A few cultivars (domesticated or human modified) species were present at the site. This included *Hordeum vulgare* (barley) and *Avena* (oat). *Hordeum vulgare* was also found in the topsoil. Some edible berries were also detected in the topsoil and in the cultural layers including *Rubus idaeus* (red raspberry), *Empetrum nigrum* (crowberry), *Vaccinium myrtillus* (European blueberry), *Vaccinium uliginosum* (bog bilberry) and *Vaccinium vitis-idaea* (lingonberry).

Charred seeds from macrofossil analysis only identified four of the 123 taxa identified by sedaDNA analysis (Table 2). Charcoal samples used for radiocarbon dating additionally identified tree species found in the sedaDNA (Cerbing, 2025) (Table 2). However, the results between methods are not directly comparable as macrofossil, charcoal and sedaDNA samples were not taken from the same location, with the exception of cooking pit 340. Macrofossil analysis from this feature identified only two unidentified charred seeds.

**Table 2.** Multiproxy comparison of plant results.

| SedaDNA ID | Taxonomic level | Macrofossil/ charcoal ID | Taxonomic level |
|---|---|---|---|
| Carex | genus | Cyperaceae/carex | genus |
| Ranunculus acris | species | Ranunculus | genus |
| Alsineae | tribe | Stellaria media | species |
| Rumex | genus | Rumex acetosella | species |
| Alnus incana | species | Alnus | genus |
| Betula | genus | Betula | genus |
| Pinus | genus | Pinus | genus |
| Salix | genus | Salix/Populus | genus |
| Populus tremula | species | Salix/Populus | genus |
| Sorbus aucuparia | species | Sorbus/Prunus | genus |
| Prunus padus | species | Sorbus/Prunus | genus |

The initial ordination from the PCA analysis resulted in three samples grouped distinctly from the other samples, Topsoil, P15308, and P14136 (Figure 3 and S25). There were six taxa only present in these samples (*Aconitum lycoctonum*, *Delphinium elatum*, *Calluna vulgaris, Plantago major*, *Humulus lupulus*, *Larix sibirica*). *Calluna vulgaris*, however, is not found in the topsoil. Some features, including cooking pits 270, 340 and hearth feature 11450 show some distinctions most likely due to the abundance of dwarf shrubs in the cooking pits compared to the other features and the presence of taxa only present in hearth 11450.

**Figure 3:**
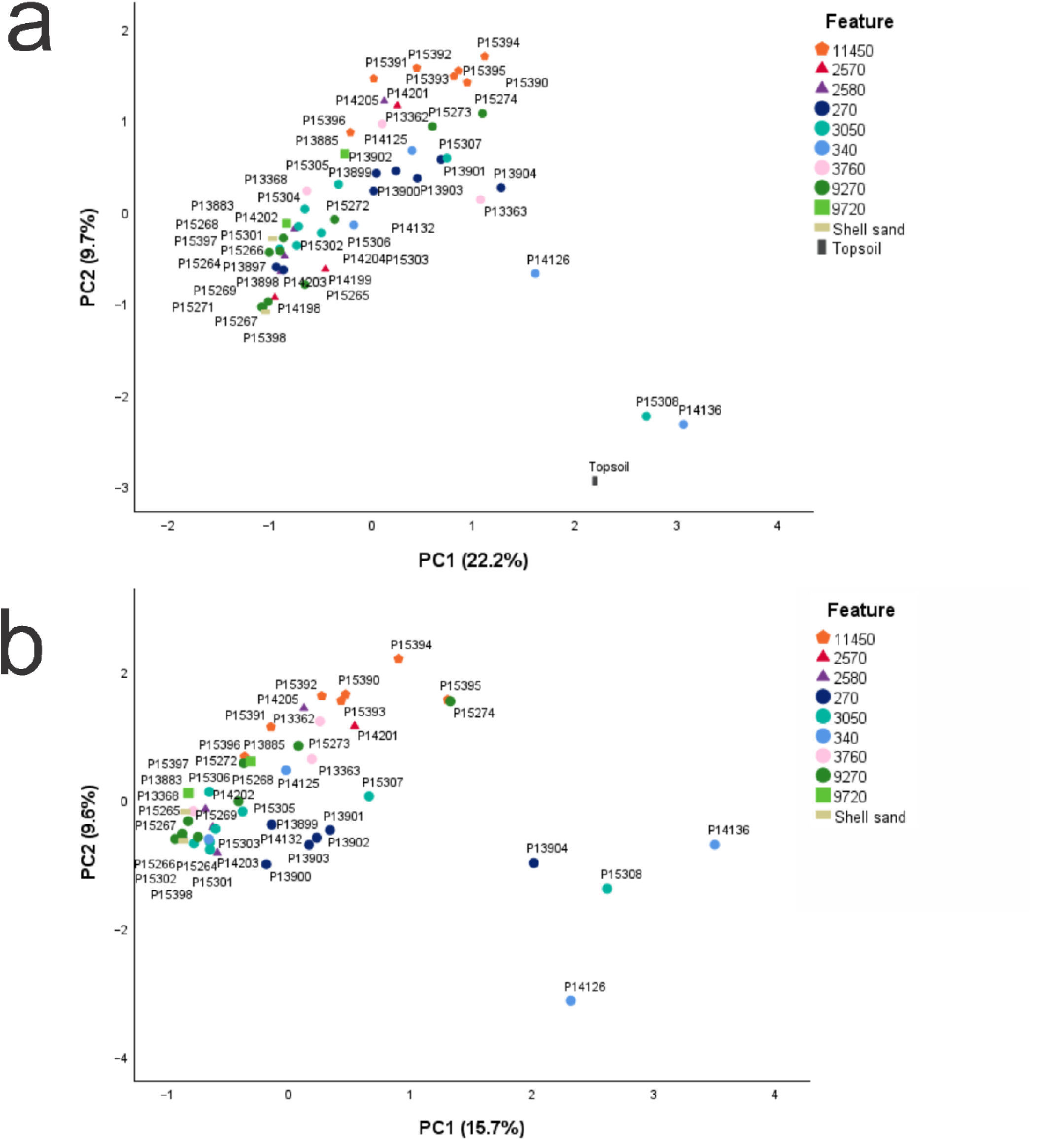
PCA analysis of samples with and without species present in the topsoil. Colors of sample plots represent feature and symbol shape represents feature type with **a)** plotting all plant taxa present and **b)** plotting only plant taxa found in the archaeological features but not in the topsoil.

When the topsoil species were removed from the analysis, 53 plant taxa remained. No taxa remained in samples P14198 and P14199 from fire pit feature 2570 and sample P15271 from feature 9270. The majority of tree and dwarf shrub species were removed with only *Quercus*, *Calluna vulgaris*, and *Arctous alpina* remaining. The species that remain are mainly forbs and graminoids (44 of 53) that grow in moist environments, meadows, and pastures (Vorren, 1986; Elven et al., 2022) (Supplementary data 12) (Figure S25). Samples from the top of features 270, 340, 9270, and 3050 (P13904, P14136, P15308, and P15274) were plotting separately from the rest of the samples suggesting differences in species composition by depth. However, samples P14126 from feature 340 are also plotting with those samples suggesting similarity in species composition throughout this feature. For the hearth and cooking pit features, samples within the same feature are clustering together, with the exception of the above-mentioned samples. While there is decreased variability after removing the topsoil species, samples within the hearth and cooking pits continue to cluster (Figure 3).

PCoA and PERMANOVA analyses were performed on the four features with more than five samples, cooking pit features 270, 3050, 9270 and hearth feature 11450 (Figure S26). The analysis was first run with species in the topsoil and then without species found in the topsoil. The PERMANOVA results remain significant after removing topsoil species from the analysis. Both PERMANOVA analyses had significant p-values of 0.001 and 0.0006 respectively, indicating the variation of species composition was greater between features than within features both with and without topsoil species (Figure S26).

### 3.2 Animal results

Across the 52 successful samples, 10 mammals, eight fish/aquatic species, two birds, and 13 annelids (worms) were detected (Figure 3). In total, 33 animal taxa were detected (Figure S20-S23, supplementary data 11). Three taxa were found in the extraction controls that were also found in the samples but are known as common contaminants (*Felis catus*, *Gallus*, and *Sus scrofa*). The sequences did not occur in more than one repeat of the controls. To account for this potential contamination, any occurrence of these DNA sequences in less than two repeats of a sample were removed from analysis. Based on number of repeats, the main mammalian taxa present in all features were domesticates including *Bos taurus* (cow), *Ovis* (sheep), *Capra* (goat), and *Sus scrofa* (pig). These species were also found in the topsoil samples. Additional species found in both the archaeological contexts and topsoil include *Microtus agrestis* (short-tailed field vole), *Lemmus lemmus* (Norway lemming), *Clupea harengus* (Atlantic herring), *Gadidae* (cod), *Pollachius virens* (pollock), and *Lumbricus rubellus* (redhead worm). *Equus* (horse) is only present in the archaeological features with five repeats in total. Three bird taxa were detected, *Lagopus lagopus* (willow ptarmigan), *Corvus* (crow), and *Larus* (gulls). A variety of fish species were also detected in feature 3050 (sample P15308) and include *Clupea harengus*, *Gadidae*, *Pollachius virens, Melanogrammus aeglefinus* (haddock), and *Pleuronectidae* (flounder). *Salmo salar* (Atlantic salmon) is also found in feature 3760 (sample P13363).

The taxa identified by both zooarchaeology and sedaDNA include *Ovis*, *Capra*, *Sus scrofa*, *Bos taurus*, *Gadidae, Pollachius virens,* and *Larus* (identified to *Larus argentatus* by zooarchaeological analysis). *Ovis* and *Capra* comprised 38.7% of the total mammal bones identified with *Bos taurus* contributing 4.7% and *Sus scrofa* 0.6%. *Gadidae* comprised 50-80% of identified fish remains (Cerbing, 2025). Additionally, macrofossil analysis identified a large proportion of burnt, unburnt, and fragmented bone material including a high number of fish bones compared to those identified in the faunal remains.

Annelids (worms) were also found across all features. In total, 36 samples contained annelids, including one cold adapted species (*Hemlea perpusilla*). All the worms are non-burrowing (eoigeic) enchytraeids except for the species, *Lumbricus rubellus*, which is a burrowing (anecic) earthworm. Feature 270 contained the highest amount of annelid DNA with 117 repeats and five species present. The fire pit features 2570 and 2580 contained the least amount of annelid DNA with only three repeats across two species present in the former and four repeats and only one species in the latter. The results from the co-occurrence analysis found multiple strong correlations between the presence of DNA from domestic taxa and annelids compared to other taxa, specifically with the annelid species *Lumbricus rubellus* and genus *Aporrectodea* (Figure 5). The strongest co-occurrence was found between *Bos taurus* and *Lumbricus rubellus* (p-value<0.00001). Other taxa were found to significantly co-occur with *Lumbricus rubellus*, including *Ovis, Aporrectodea*, *Capra*, *Sus scrofa* and *Gadidea* (p-values = 0.00011, 0.00010, 0.00123, 0.00158 and 0.03987 respectively). All significant co-occurrences were positive associations.

**Figure 4.**
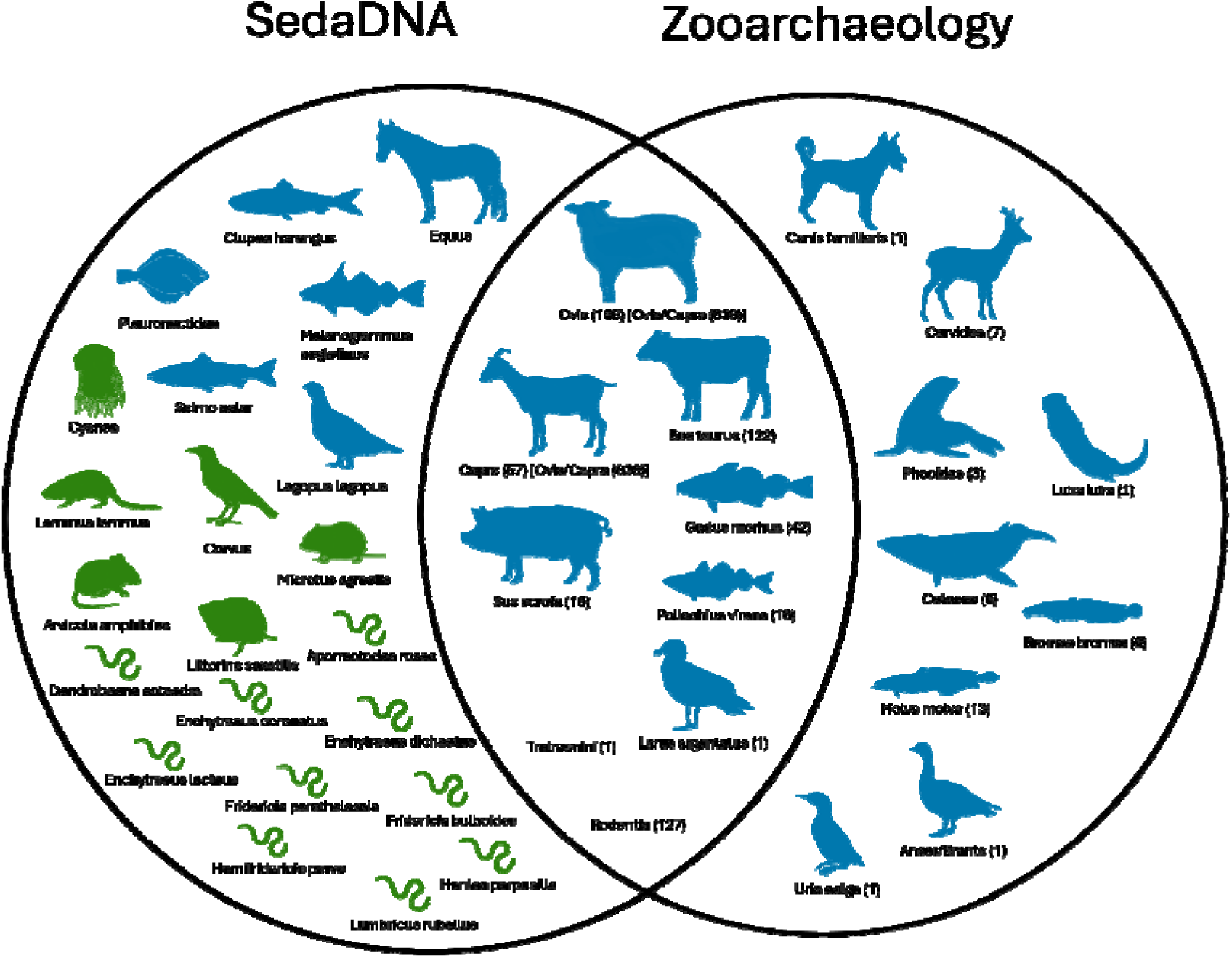
Overlap of species detection between sedaDNA and zooarchaeology analysis. Species in blue were consumed or likely to have been consumed at the site and species in green are found naturally in the surrounding environment or were unintentionally introduced. Species in opaque colors identified to a higher taxonomic level in the faunal remains but most likely match to species identified in sedaDNA. Number of Identified Specimens (NISP) values are included for species found in the faunal assemblage. Species only found in the topsoil layers were not included.

**Figure 5:**
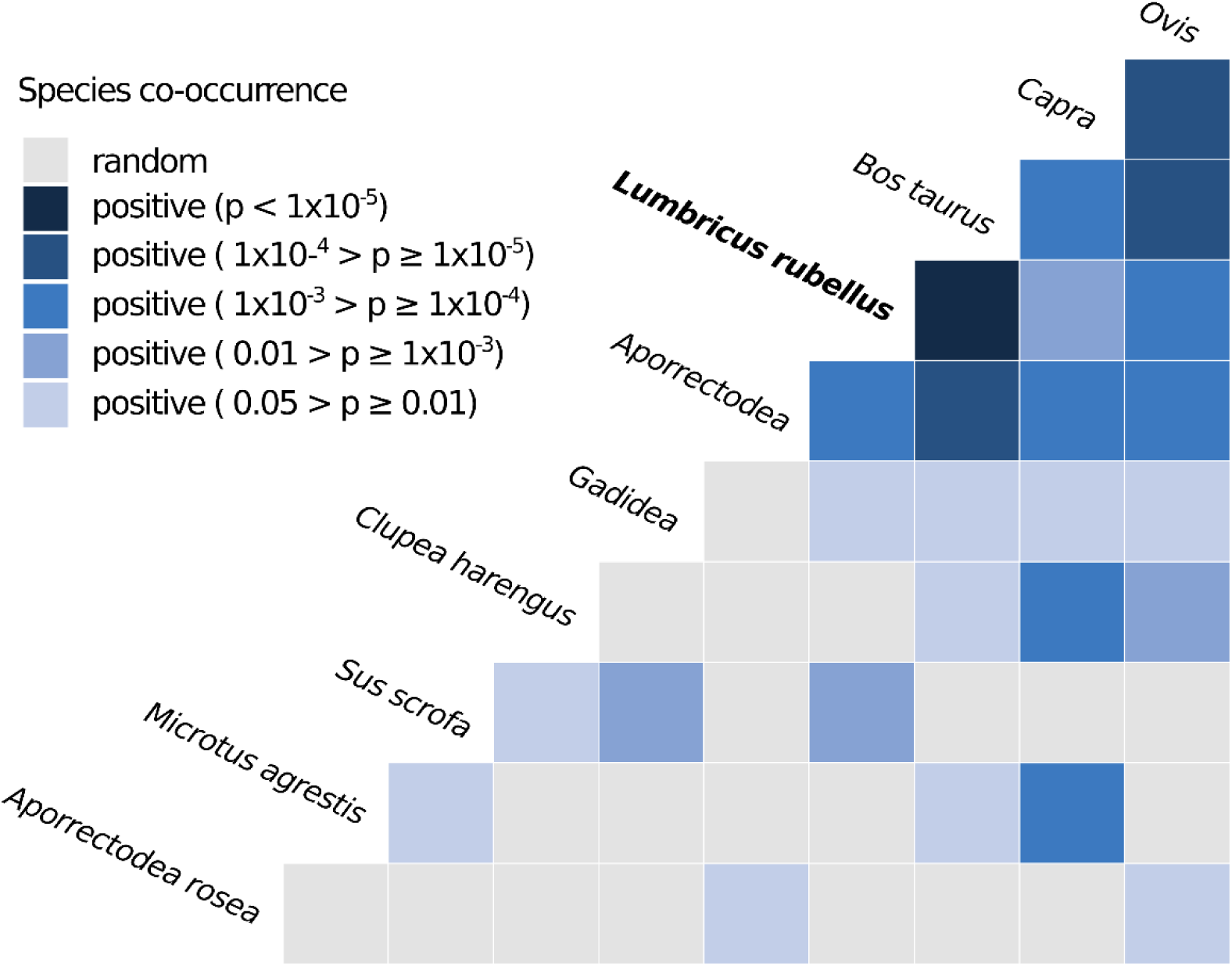
Co-occurrence matrix demonstrating the co-occurrence of domestic taxa with annelids, particularly *Lumbricus rubellus*. A positive result indicates a significant association between taxa. A random result indicates no or an insignificant association between taxa.

## 4. Discussion

This study successfully sequenced sedaDNA from multiple heated features at the Iron Age site of Rødskjær. The plants present at the site come from a wide variety of ecological zones and would not be found together in the natural environment (Elven et al., 2022). Additionally, many of the samples were species-rich compared to what might be expected in natural soils and sediments (Ariza et al., 2022). Both findings suggest anthropogenic activity at the site (Chen et al., 2026). The number of shared plant and animal species between the topsoil samples and archaeological sediments made it difficult to interpret if these species were introduced during the Iron Age or from agricultural or pastoral use of the site in later time periods. While many of the species are native and could be found in Iron Age Norway, there are indications of younger sediments and topsoil mixing with the Iron Age sediments through worm bioturbation as well as indicators of root penetration in some contexts. The amount that these processes are affecting each feature varies. While all features sampled at Rødskjær are heated features, their construction, use and post-use processes are highly variable and complex (Aldeias, 2017). Heated archaeological features also often contain a mix of sediments, fire cracked rocks, charcoal and microfossils that have been variably affected by the heating process. However, some patterns emerged of how sedaDNA was being deposited and preserved in the features.

There are at least two plant species (*Delphinium elatum* and *Larix sibirica*) found in the shallow sediment samples from the features that are plants introduced to Norway after 1700 AD (Elven et al., 2022). The presence of these species in samples P15308, P14136, and P14201 suggests plant roots penetrating the sediments or intermixing with modern soil. Additionally, the presence of the deep burrowing (anecic) worm species *Lumbricus rubellus* was found to significantly co-occur with that of domestic mammals within the cooking pit and hearth features (Figure 4). Earthworms (anecic and endogeic) can move archaeological materials vertically and horizontally through a sediment profile and burrowing worm species including *Lumbricus rubellus* can transport topsoil up to three meters deep into sediment (Escosteguy and Fernández 2017; Jerand et al., 2023). This suggests that there is a certain amount of bioturbation occurring where burrowing worm species are moving younger soils and sediments into the features. However, the lack of these modern garden taxa deeper into the sediment profiles suggests topsoils are not fully mixing with the archaeological sediments. There are additionally some indications of spatial differences in species composition between features that could be explained by the residential use of the site. The fire pit features 2570 and 2580 do not contain *Lumbricus rubellus* DNA but do contain DNA from two *Enchytraeus* species commonly known as pot worms, that are shallow dwelling species often found in compost and organic rich soils (Amossé et al., 2016). It was not expected to find these species in these features as they are comprised of alternating shell-sand and charcoal layers. However, these two features were found beneath a modern garden and *Enchytraeus* was likely introduced from the above garden soils (Figure S2).

The high number of species present in the cooking pit and hearth features compared to the fireplace and fire pits suggests the sedaDNA preserved in the cooking pits and hearth is from the more organic rich backfill sediments present in these feature types. Backfill sediments introduced soon after the disuse of each feature would be representing species present at the site within a similar timeframe of the feature. The results of the PCA analysis showed some difference in the species composition between the samples from the top of the features compared to samples from deeper in the features for cooking pits 270, 9270, and 3050. This additionally indicates the mixing of sediments at the top of the features but not further into the features. Feature 340 did not follow this pattern suggesting similarity in species composition throughout this feature. This feature was not dated and most samples from this feature did not yield sedaDNA so it is difficult to interpret why this may be. However, overall the PCoA found more variability in species composition between features than between samples within the same feature. This indicates species composition differences between features due to spatial, temporal, or use differences within the features. The PCoA analysis showed the most differences in plant species composition between the hearth feature 11450 and the cooking pits, potentially indicating different use of this feature (Figure S26). We could not fully assess this due to not all the features being dated and a limited amount of topsoil samples taken. A systematic sampling of topsoil from across the site area would be beneficial to assess spatial differences in species composition from across the site. This could not be done at Rødskjær due to the site being surface stripped before sedaDNA samples were taken, but should be conducted at other sites when conducting analysis of sedaDNA samples. A vegetation survey would also assist in the evaluation of plant species present across the site area at the time of excavation.

To better evaluate the contribution of sedaDNA coming from later time periods, a record of the post-site history is needed. While the use of the site after the Iron Age is not fully known, the finds at the site indicate some activity in the Viking Age (Cerbing, 2025). There is no archaeological evidence of permanent settlement during this time period from the site, but Rødskjær is also near (600m to the north) the site of Sandtorg, which is known from archaeological and historical evidence to have been used as a farm in the medieval period and is a suspected trading site from the Viking Age through to the medieval period. This makes it likely that Rødskjær had near continuous activity after the Iron Age as agricultural or pastureland as the plant species not found in the topsoil also suggest (Lange et al., 1873; Krokmyrdal, 2021).

Comparing the faunal remains, macrofossil, and charcoal analysis to the sedaDNA confirms the Iron Age presence of some plant and animal species found both in the topsoil and archaeological sediments. Comparing findings between the sedaDNA and the faunal remains showed relatively low overlap (Supplementary data 14). However, the most abundant taxa at the site were identified by both methods (Figure 4). The taxa that were only identified by zooarchaeological analysis were in low abundance at the site with each species being only identified by the presence of one to three bone elements. SedaDNA is less likely to detect rare taxa unless sediment samples are taken directly from where the taxa were present or directly deposited after death (Edwards et al., 2018; Murchie et al., 2023). The co-occurrence of *Lumbricus rubellus* with the domesticate taxa indicates these annelids transporting DNA of these species from the topsoils to the archaeological sediments. However, the faunal remains indicate that domesticate taxa, among other taxa, were being butchered on site (Cerbing, 2025). It is therefore highly likely that DNA from bone and soft tissues are being preserved in the sediment from the butchery and cooking process (Thomas et al., 2019). While three fish taxa were also found in the topsoil, the faunal assemblage confirmed the presence of cod and pollack at the site. The majority of fish sedaDNA was found in cooking pit feature 3050. The faunal assemblage, macrofossil analysis and sedaDNA all suggest an increase in fish dating from the later use period of the site. Identification of charred plant remains used for ^14^C dating identified tree species also found in the topsoil and features presumably brought in for fuel and not naturally growing on site as no plant root penetration from these species were found. Macrofossil analysis additionally confirmed the presence of four plant taxa and fragmented fish bones (Table 2). Macrofossil analysis found few indicators of trees, shrubs, or heath plants (Cerbing, 2025). Macrofossils were mostly comprised of species that grow in open areas and are indicative of grazing. For sedaDNA samples, the plant composition of species not found in the topsoil is also indicative of open pasture.

There were some taxa detected in the sedaDNA that were not detected in the topsoil or natural sediments and that were likely present during the Iron Age occupation of the site. These include *Avena* (oat), *Quercus* (oak), *Equus* (horse), and *Salmo salar* (Atlantic salmon). The presence of horse could indicate the use of these animals as transport and pack animals. The presence of horses in Iron Age Norway is confirmed by the findings of horse remains in Iron Age graves (Klokkervoll, 2015). There is also evidence from southern parts of Norway for the use of horses in ritual sacrifices in the Iron Age (Bukkemoen, 2017). While horse remains have been found at cooking pit field sites, they have not thus far been discovered at courtyard sites in Norway. The absence of horse bones at the site suggests the sedaDNA is being deposited through urine and feces (Andersen et al., 2012; Bukkemoen, 2017; Grønnesby et al., 2023). *Avena* and *Salmo salar* were also detected. *Avena* is present in four replicates across three features. It is unlikely to have been cultivated at Rødskjær but rather brought to the site. There is some evidence for oat cultivation in southern Norway during the Early Iron Age, but it is not thought to be a substantial part of the diet in this time period, and evidence for oat at archaeological sites in northern Norway is only found by two pollen studies and in low abundance (Josefsson et al., 2014; La Torre et al., 2026). As *Salmo salar* was not identified in the zooarchaeological assemblage, these fish were then most likely consumed but not a substantial part of the diet. *Quercus* is not native to northern Norway and is only sparsely grown in the region today. *Quercus* was commonly used in southern Norway in the Iron Age for making wooden objects and Iron Age bog boats made from oak have been found in northern Norway (Wickler, 2024). It is therefore likely that this species was brought to the site as wood or in the form of worked wooden objects.

## 5. Conclusion

This study has investigated the preservation of sedaDNA in heated archaeological features at the Iron Age site of Rødskjær in northern Norway. Plant and animal sedaDNA was successfully extracted from 52 of the samples including a total of 131 taxa that were not identified through macrofossil, charcoal, or zooarchaeological analysis. The study further addressed how sedaDNA integrity at the site could be affected by factors such as worm bioturbation. The number of shared taxa between the modern topsoil and archaeological sediments also posed a challenge and further sedaDNA sampling of topsoil from across the site area as well as a vegetation survey could help address this. Despite these challenges, both zooarchaeology and sedaDNA identified the most abundant taxa at the site and sedaDNA additionally identified taxa present in the features that were not identified in the zooarchaeological record. Most significantly, sedaDNA identified *Equus* (horse) which was not found in the faunal record and suggests the use of horse at the site as transport and pack animals. *Quercus* (oak), *Avena* (oat), and *Salmo salar* (salmon) were also identified and were most likely brought to the site. Additionally, an increase of fish species in the later use period of the site was observed across methods. Plant sedaDNA results identified a much greater number of plant taxa than macrofossil analysis and provided evidence for human mediated alteration of species composition at the site. With further investigation into sedaDNA integrity and depositional processes, cooking pits could provide invaluable insight into site use and environment at courtyard and cooking pit field sites above and beyond those revealed here.

## Acknowledgments

We thank Marie Føreid Merkel for laboratory assistance. We thank Anja Roth Niemi, Andreas Luneborg and the archaeology department at the Arctic University Museum of Norway for providing access to materials. Mary Lucas was funded by an internal PhD position at the Arctic University Museum of Norway.

## Data Availability Statement

The raw metabarcoding sequencing data are deposited in the European Nucleotide Archive (ENA) under the project accession code PRJEB121249. Scripts for initial processing of the metabarcoding data are at https://github.com/Y-Lammers/MergeAndFilter.

