## Supplementary figures for "Sedimentary ancient DNA analysis from the Iron Age courtyard and cooking pit site of Rødskjær in Troms, Northern Norway": Rodskjaer_Supplementary Figures_high resolution.docx

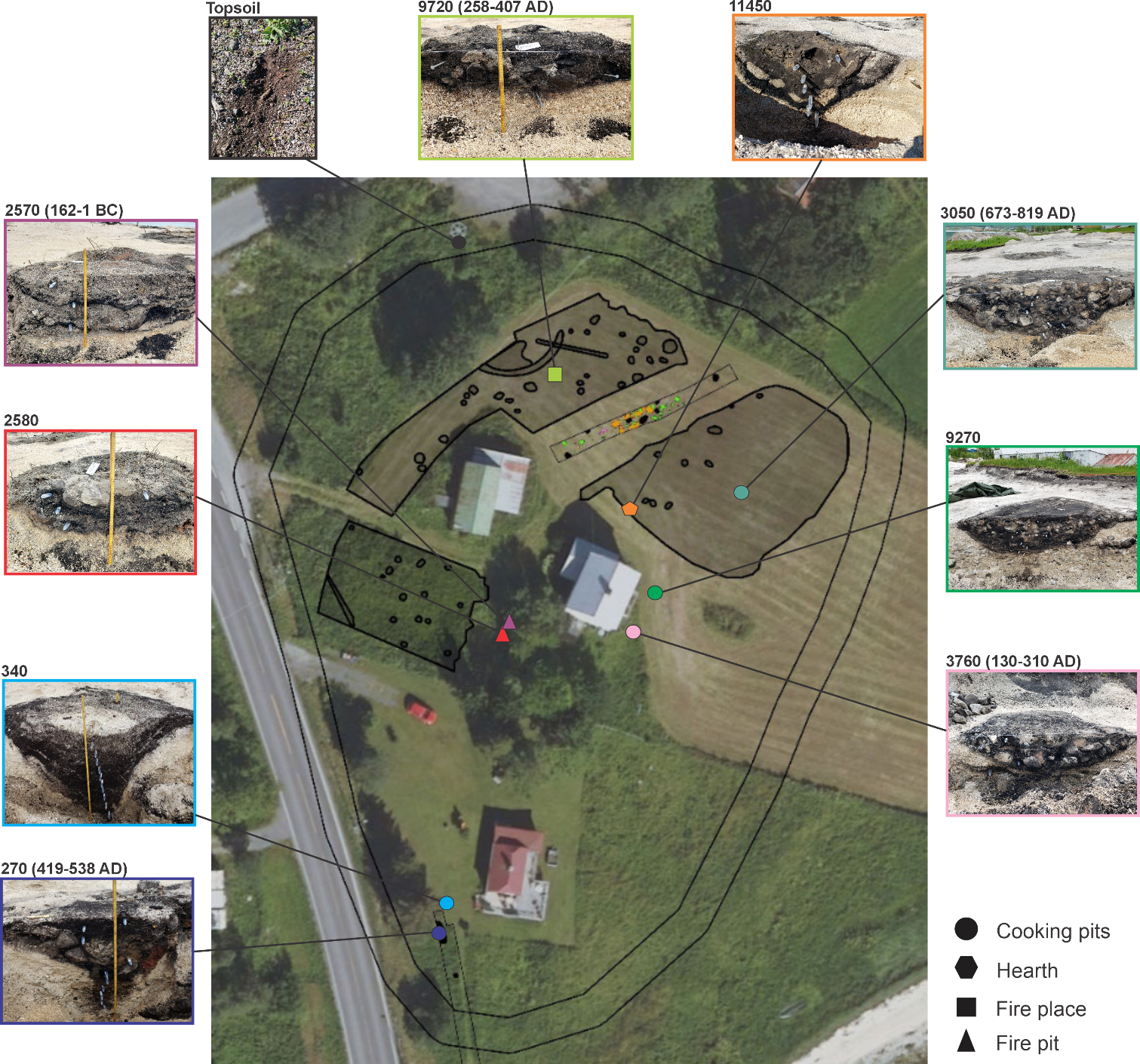

**Supplementary Figure 1.** Edited aerial photo of excavation area during first survey of the site in 2017 showing approximate locations of sampled features. Photo taken from Cerbing, 2025.

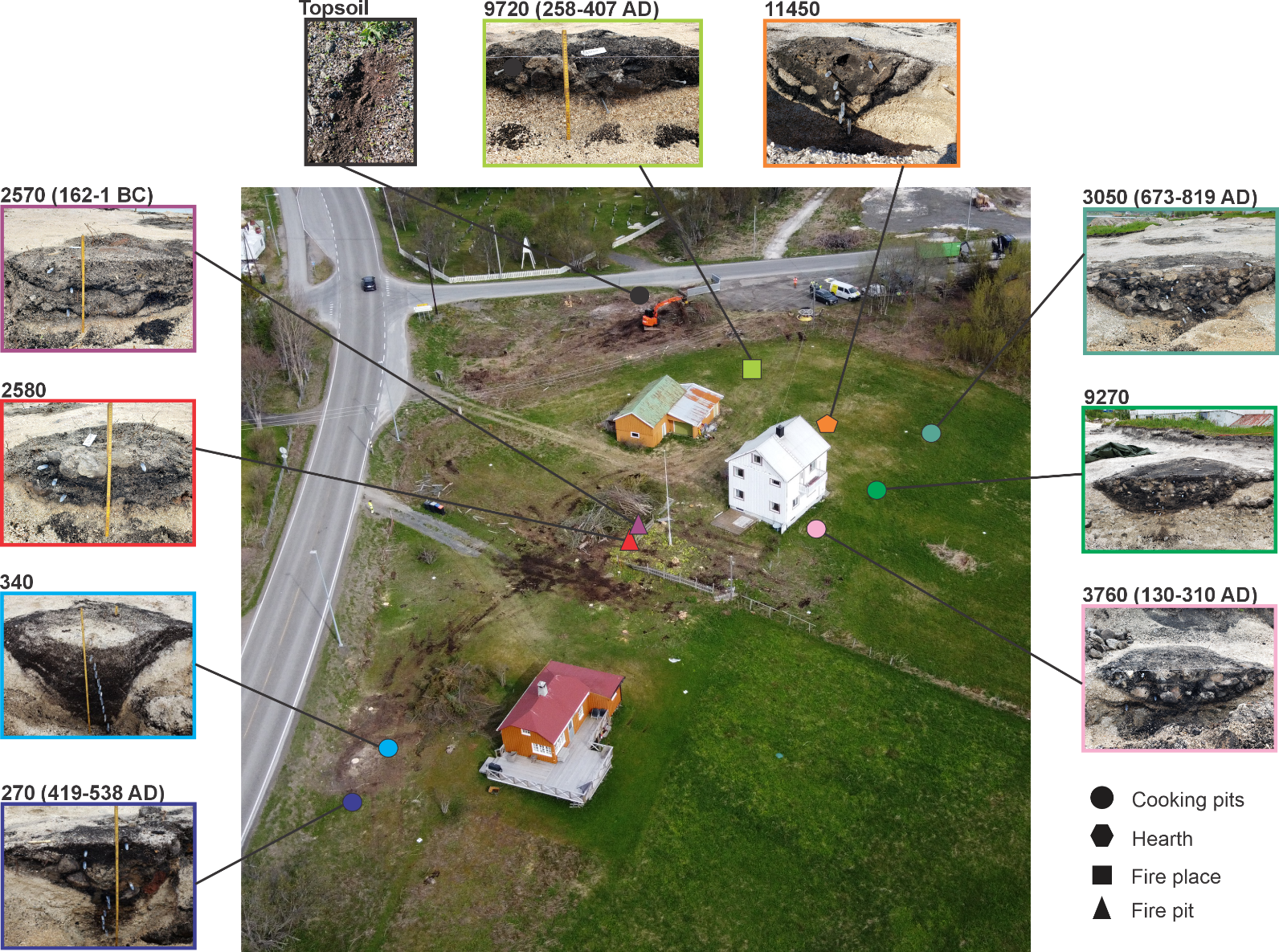

**Supplementary Figure 2.** Photo of excavation area in 2023 after clearing the site of trees and before surface stripping showing approximate locations of sampled features. Photo taken by Erik Kjellman.

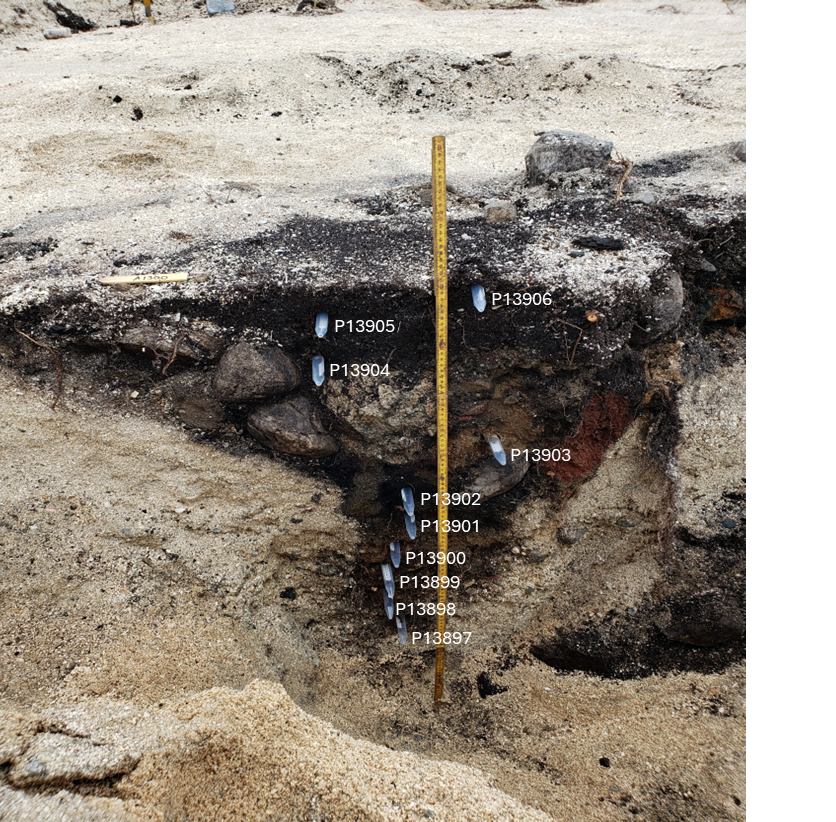

**Supplementary Figure 3.** Cooking pit feature 270 with sample labels.

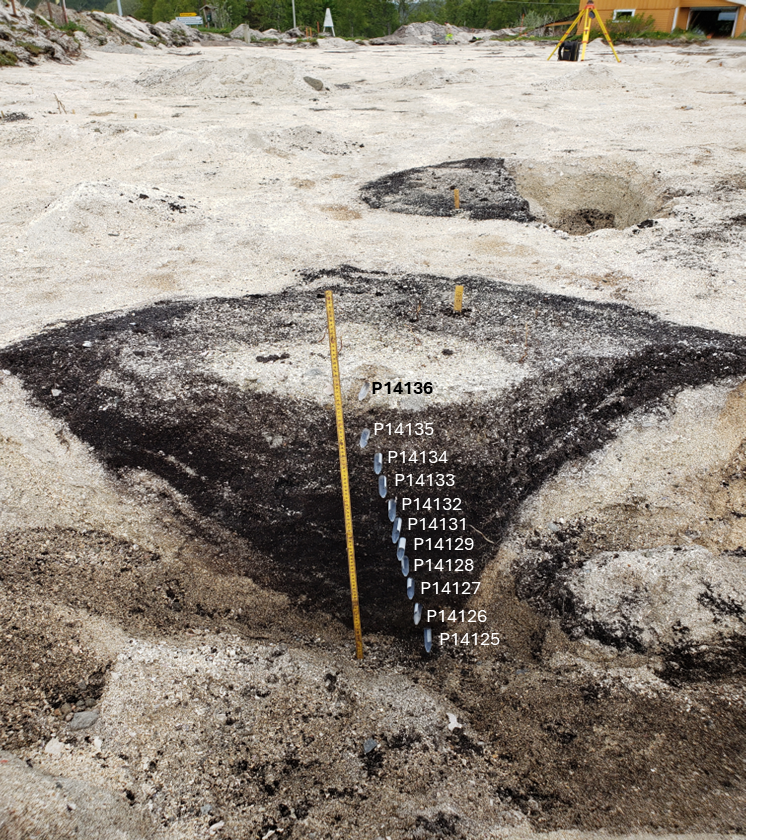

**Supplementary Figure 4.** Cooking pit feature 340 with sample labels.

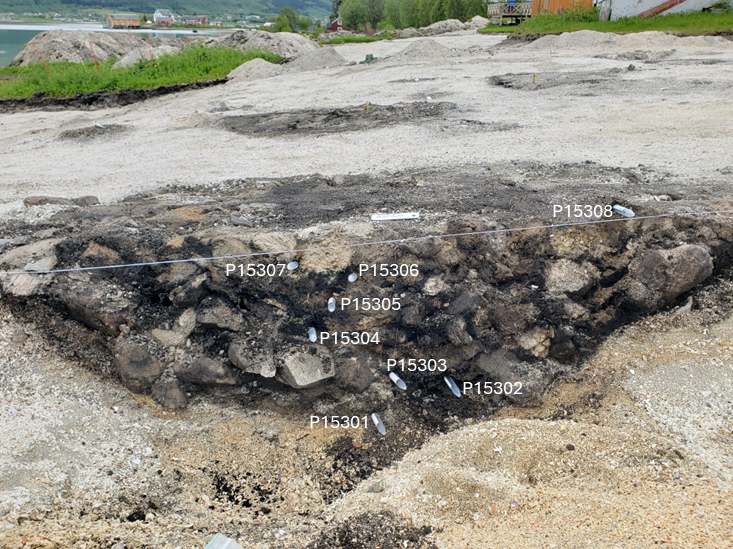

**Supplementary Figure 5.** Cooking pit feature 3050 with sample labels.

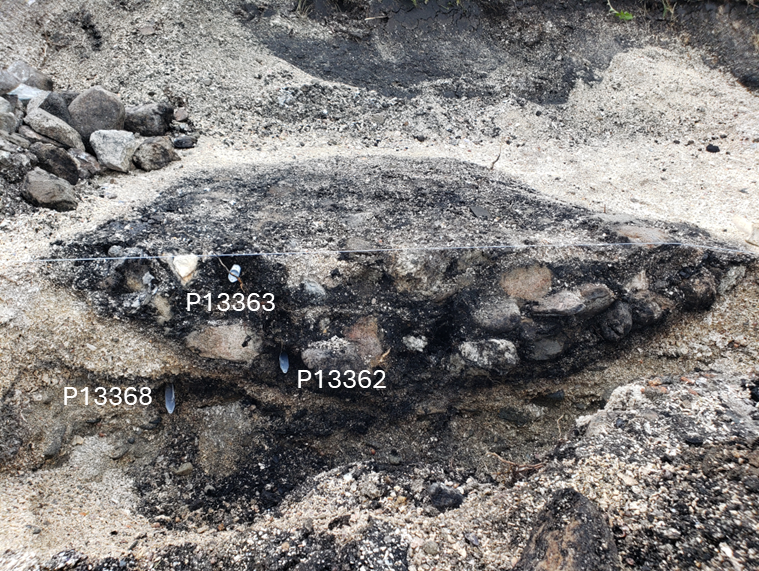

**Supplementary Figure 6.** Cooking pit feature 3760 with sample labels.

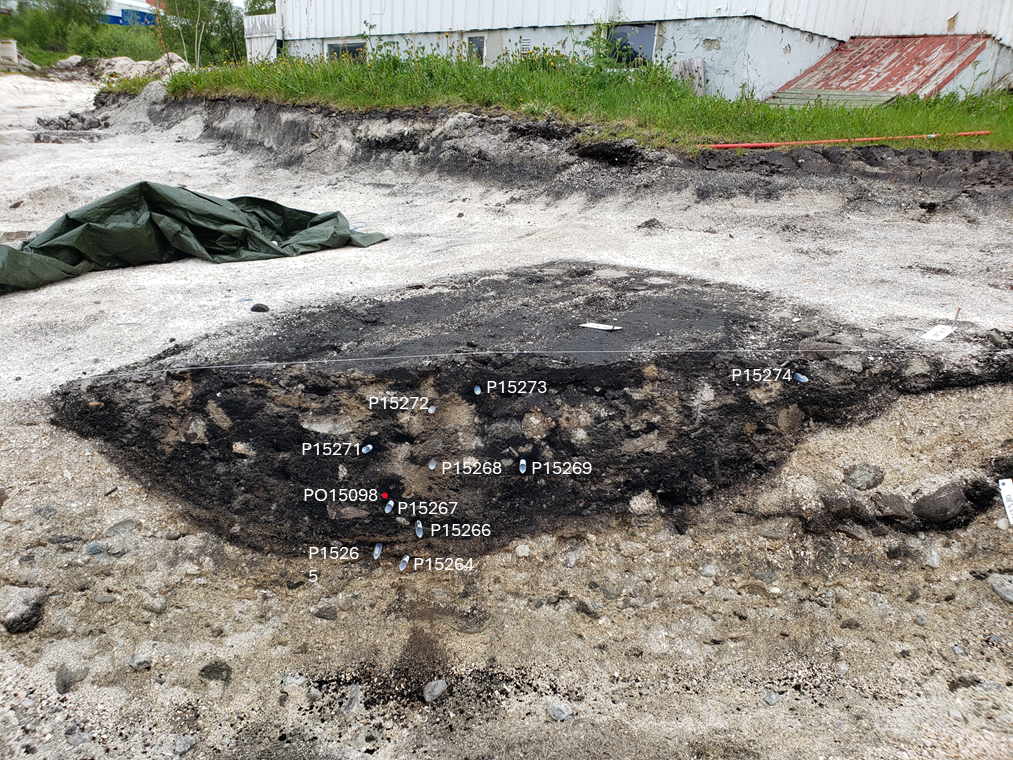

**Supplementary Figure 7.** Cooking pit feature 9270 with sample labels.

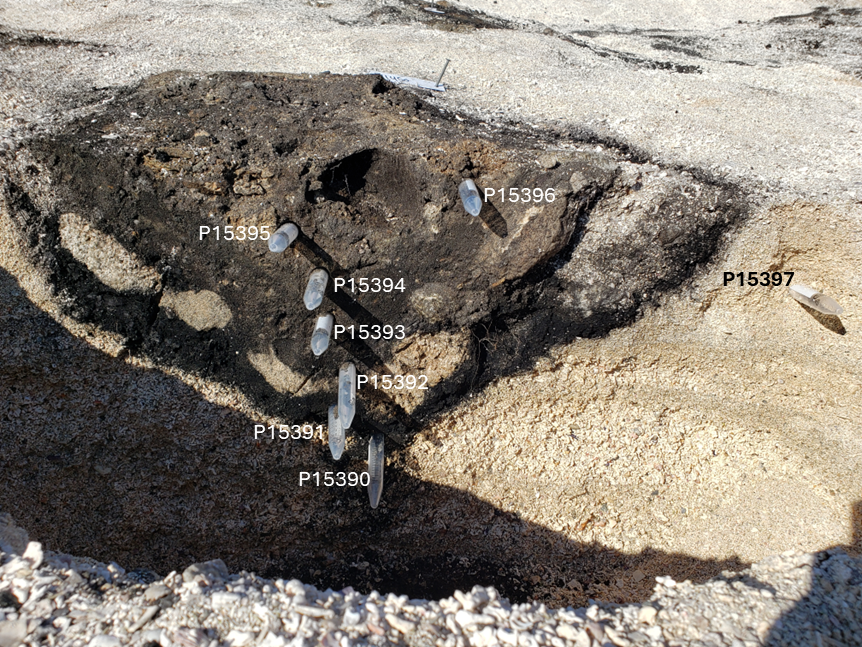

**Supplementary Figure 8.** Hearth feature 11450 with sample labels.

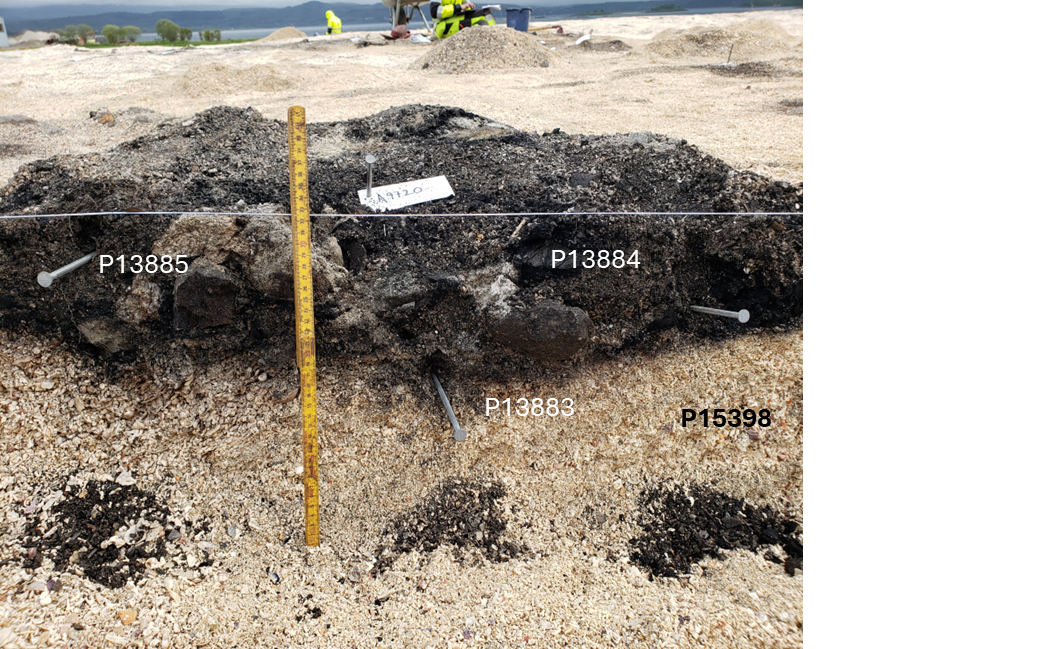

**Supplementary Figure 9.** Fireplace feature 9720 with sample labels.

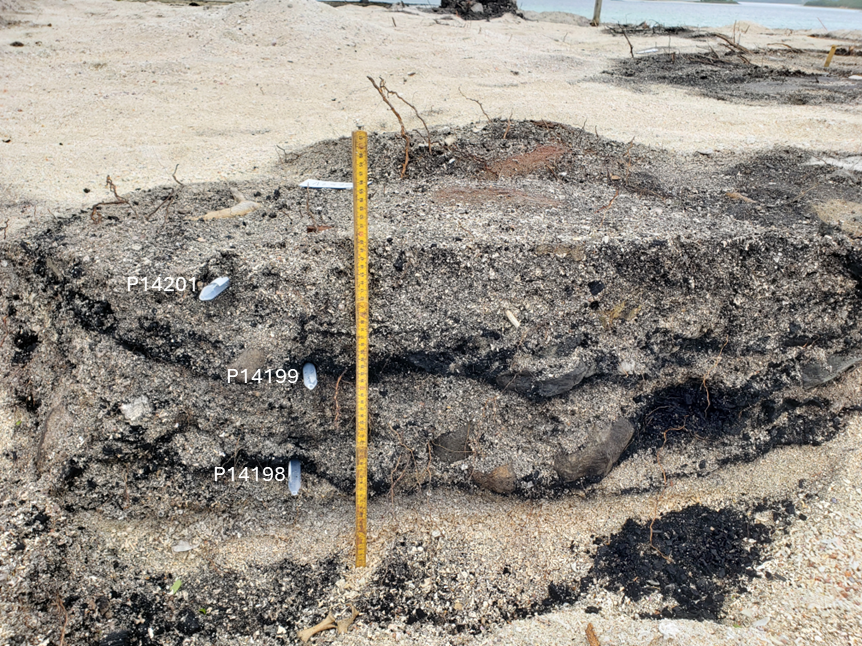

**Supplementary Figure 10.** Fire pit feature 2570 with sample labels.

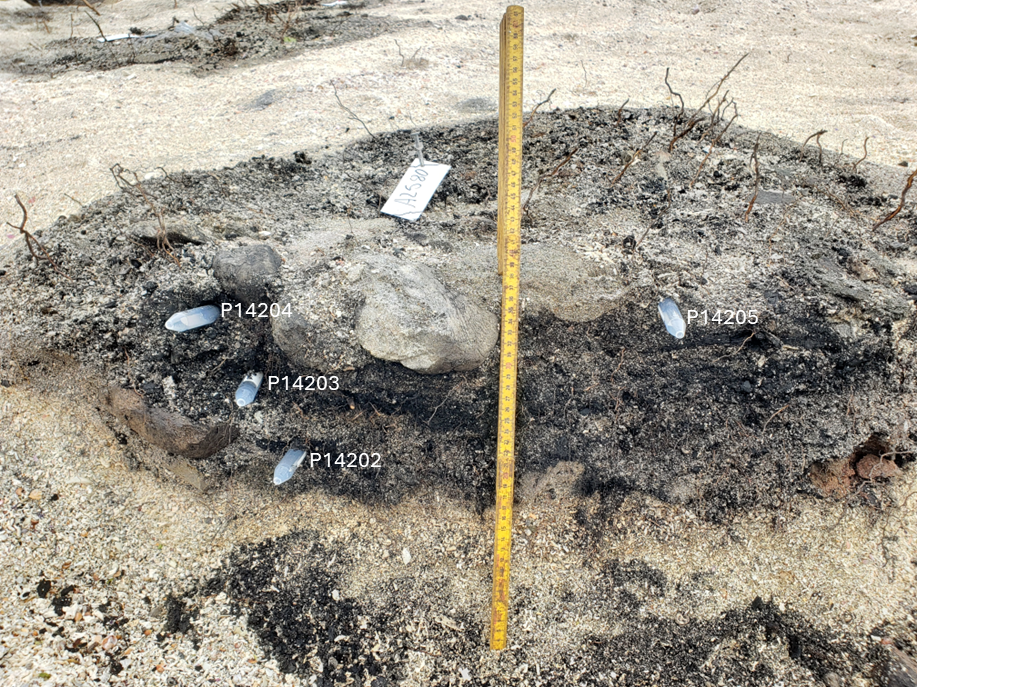

**Supplementary Figure 11.** Fire pit feature 2580 with sample labels.

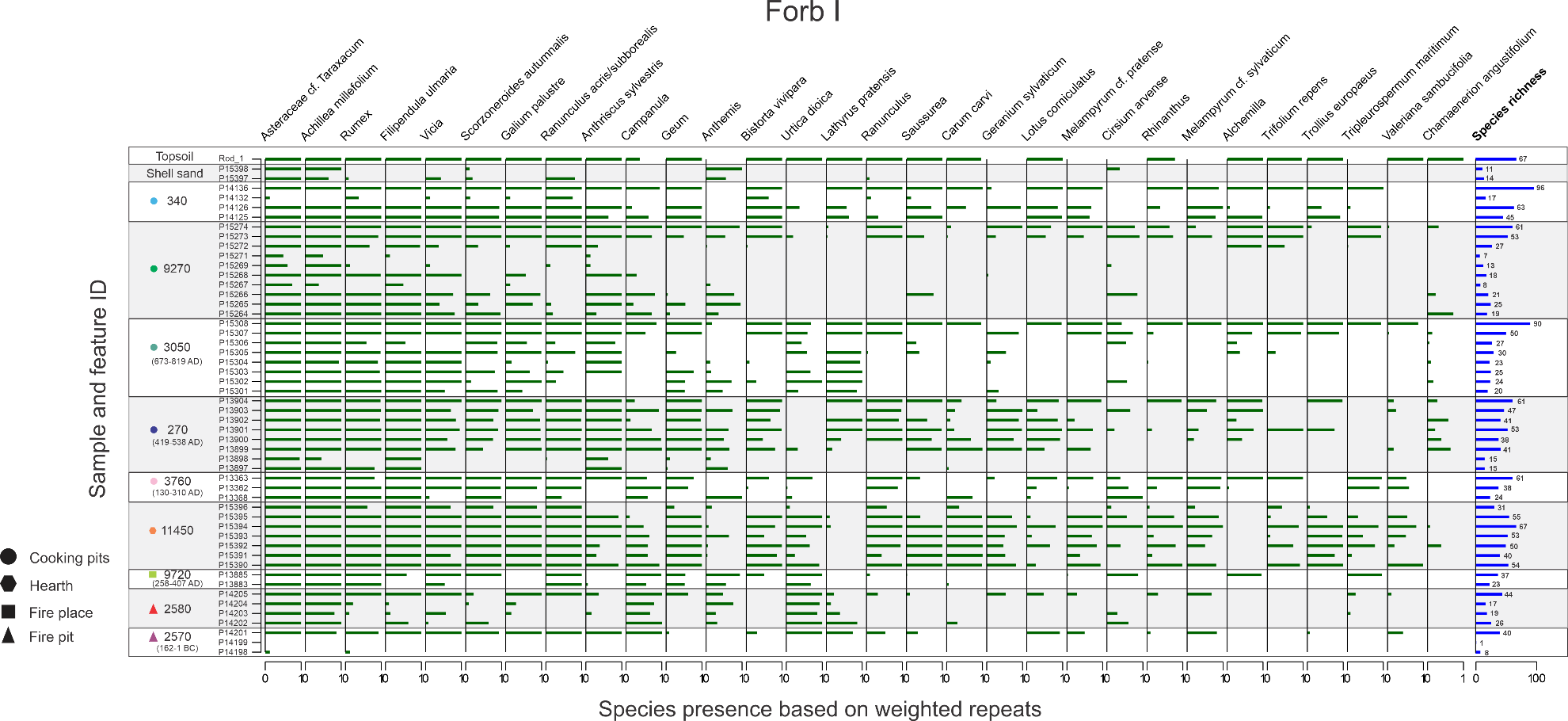

**Supplementary Figure 12.** First 30 forb species present in samples based on weighted repeats. Species present in all eight replicates of a sample equal one. Species richness for each sample represented by blue bar.

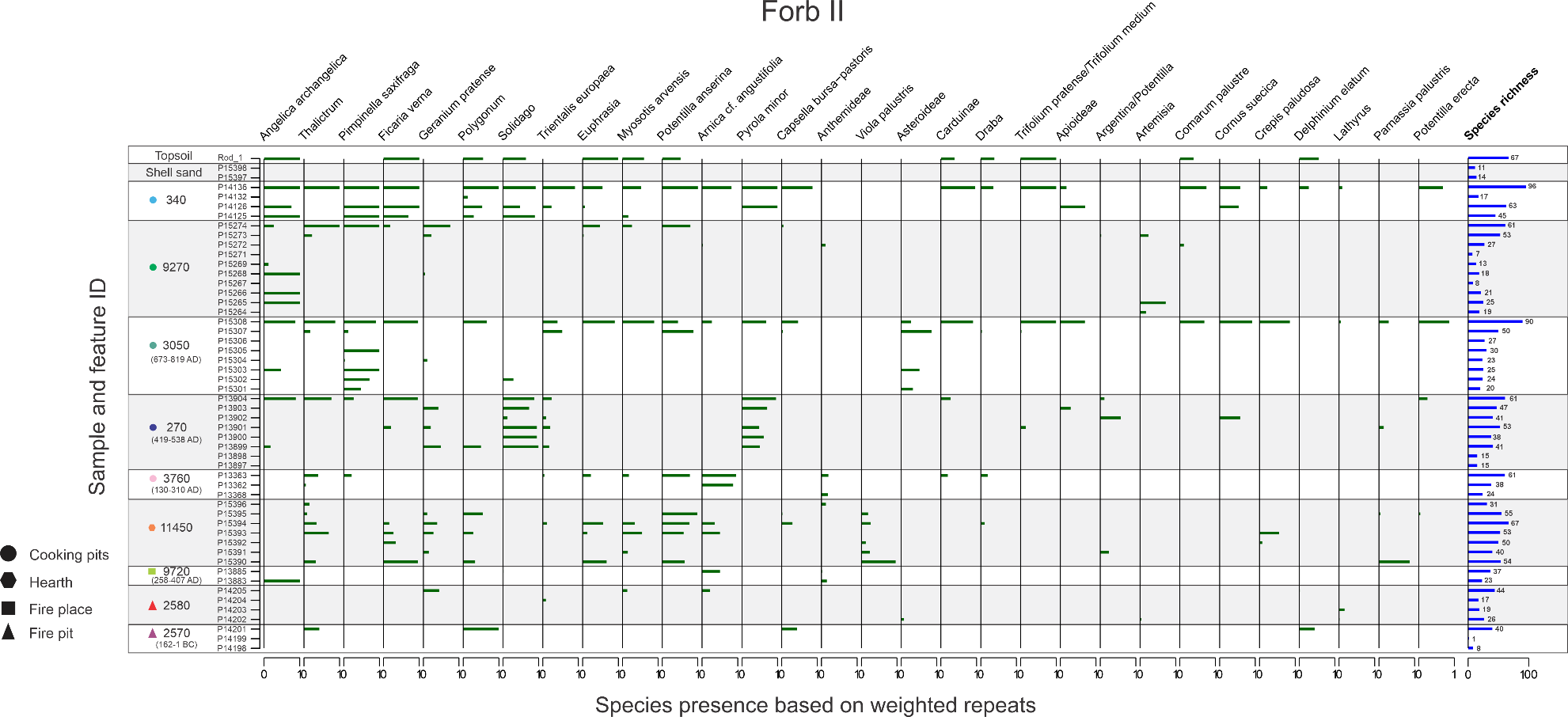

**Supplementary Figure 13.** Next 30 forb species present in samples based on weighted repeats. Species present in all eight replicates of a sample equal one. Species richness for each sample represented by blue bar.

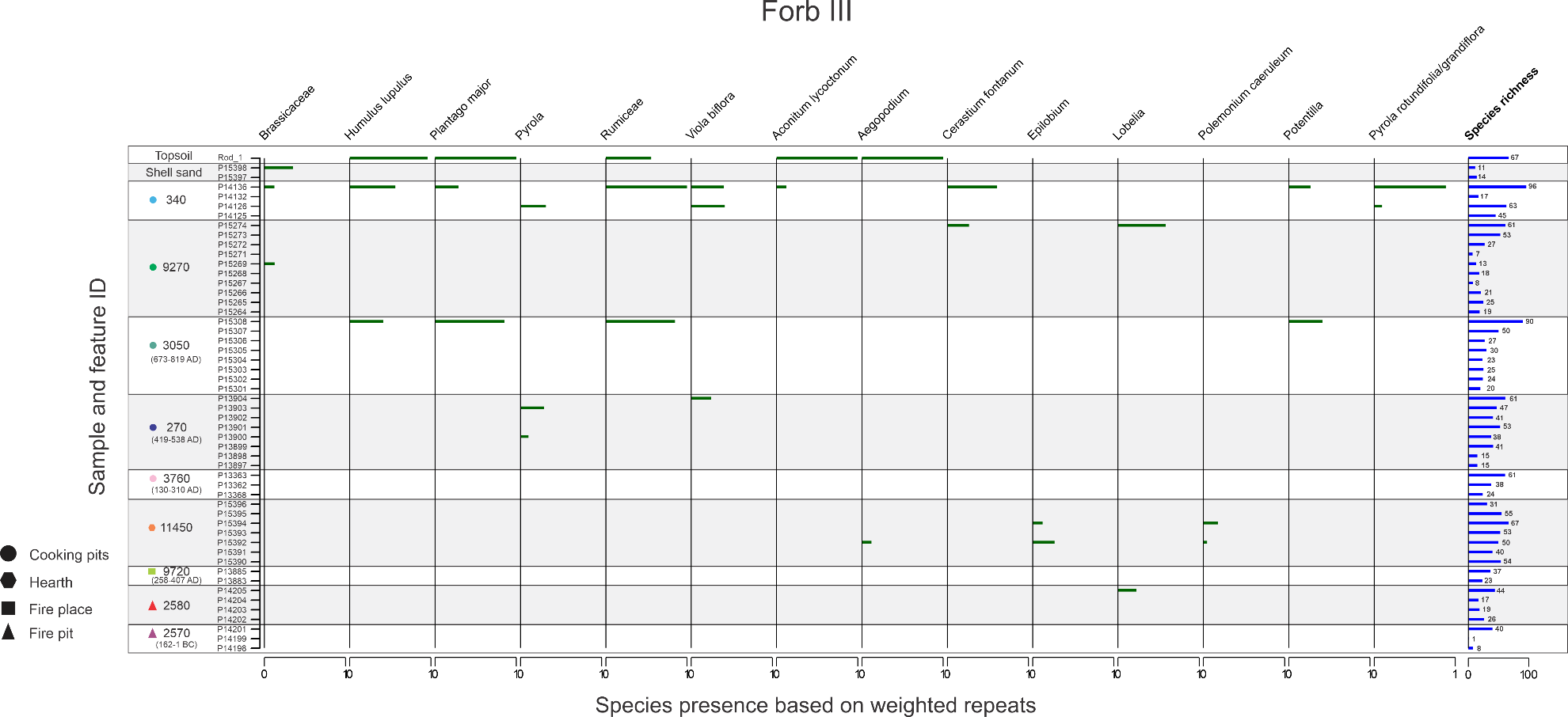

**Supplementary Figure 14.** Last 13 forb species present in samples based on weighted repeats. Species present in all eight replicates of a sample equal one. Species richness for each sample represented by blue bar.

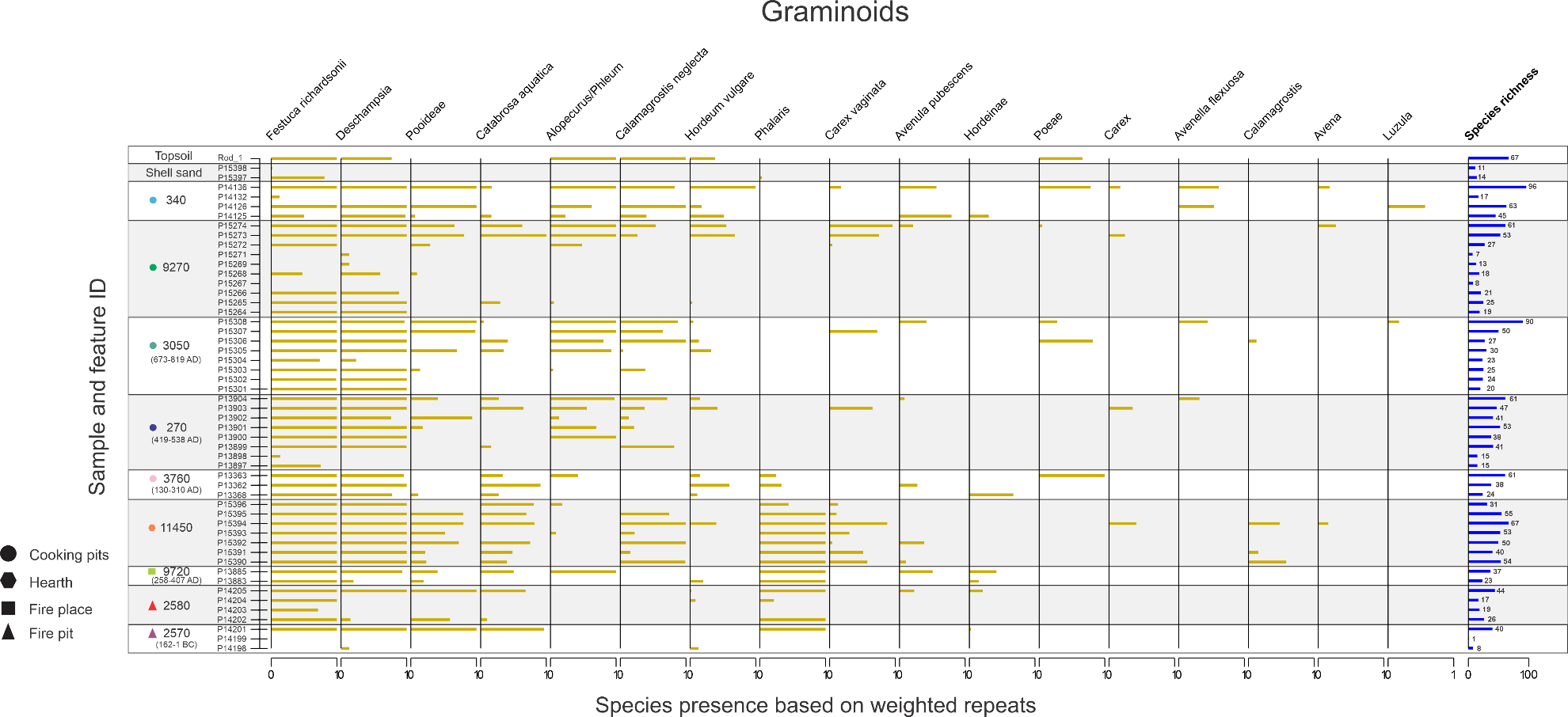

**Supplementary Figure 15.** Graminoid species present in samples based on weighted repeats. Species present in all eight replicates of a sample equal one. Species richness for each sample represented by blue bar.

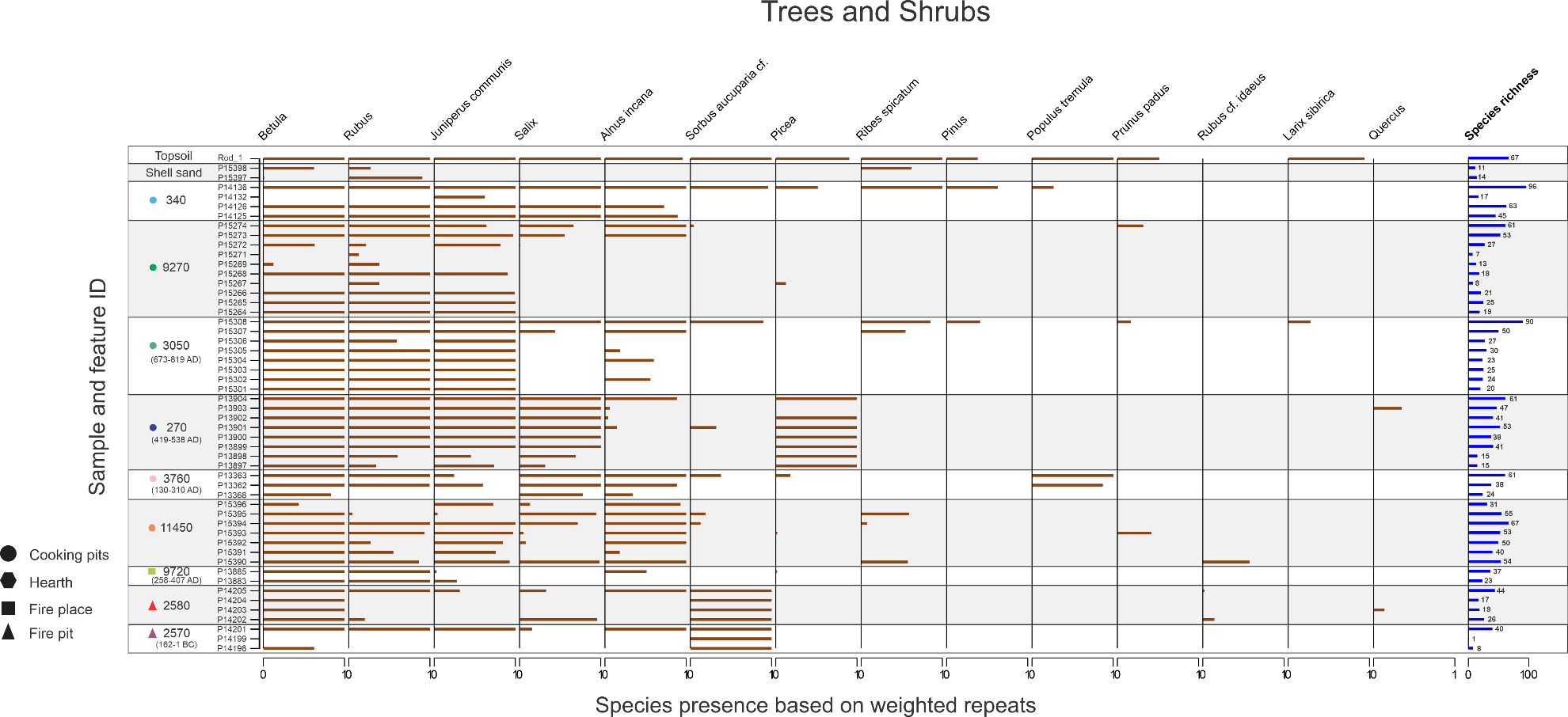

**Supplementary Figure 16.** Trees and shrub species present in samples based on weighted repeats. Species present in all eight replicates of a sample equal one. Species richness for each sample represented by blue bar.

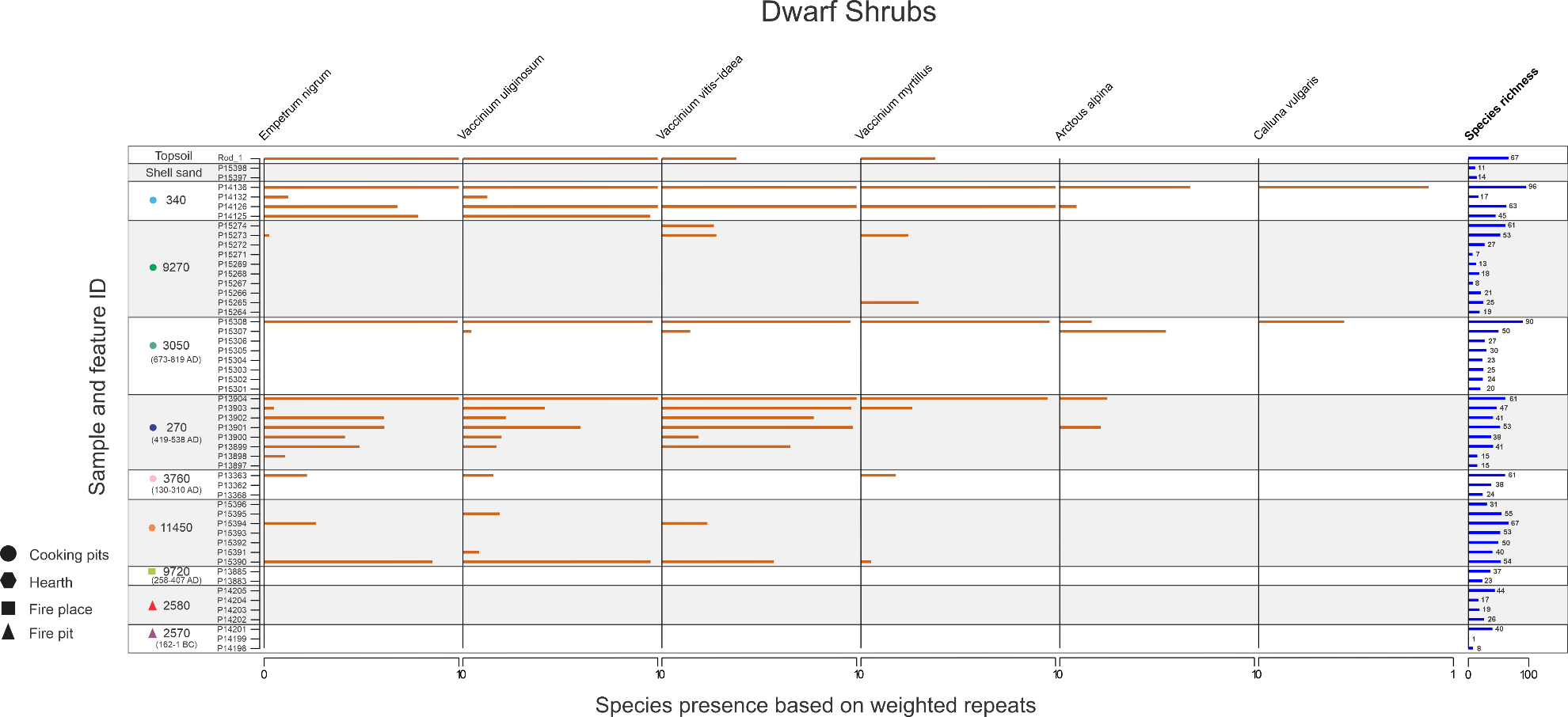

**Supplementary Figure 17.** Dwarf shrub species present in samples based on weighted repeats. Species present in all eight replicates of a sample equal one. Species richness for each sample represented by blue bar.

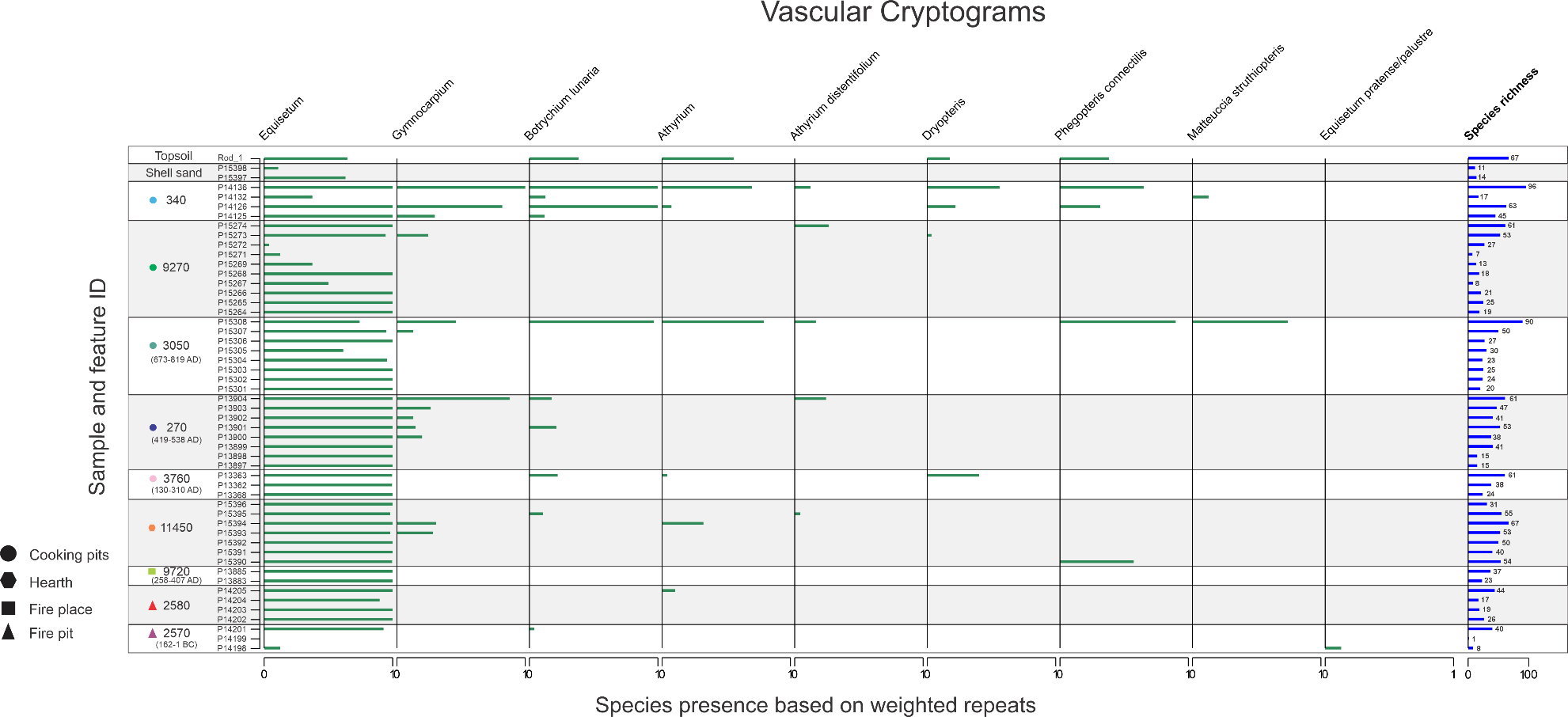

**Supplementary Figure 18.** Vascular cryptogram species present in samples based on weighted repeats. Species present in all eight replicates of a sample equal one. Species richness for each sample represented by blue bar.

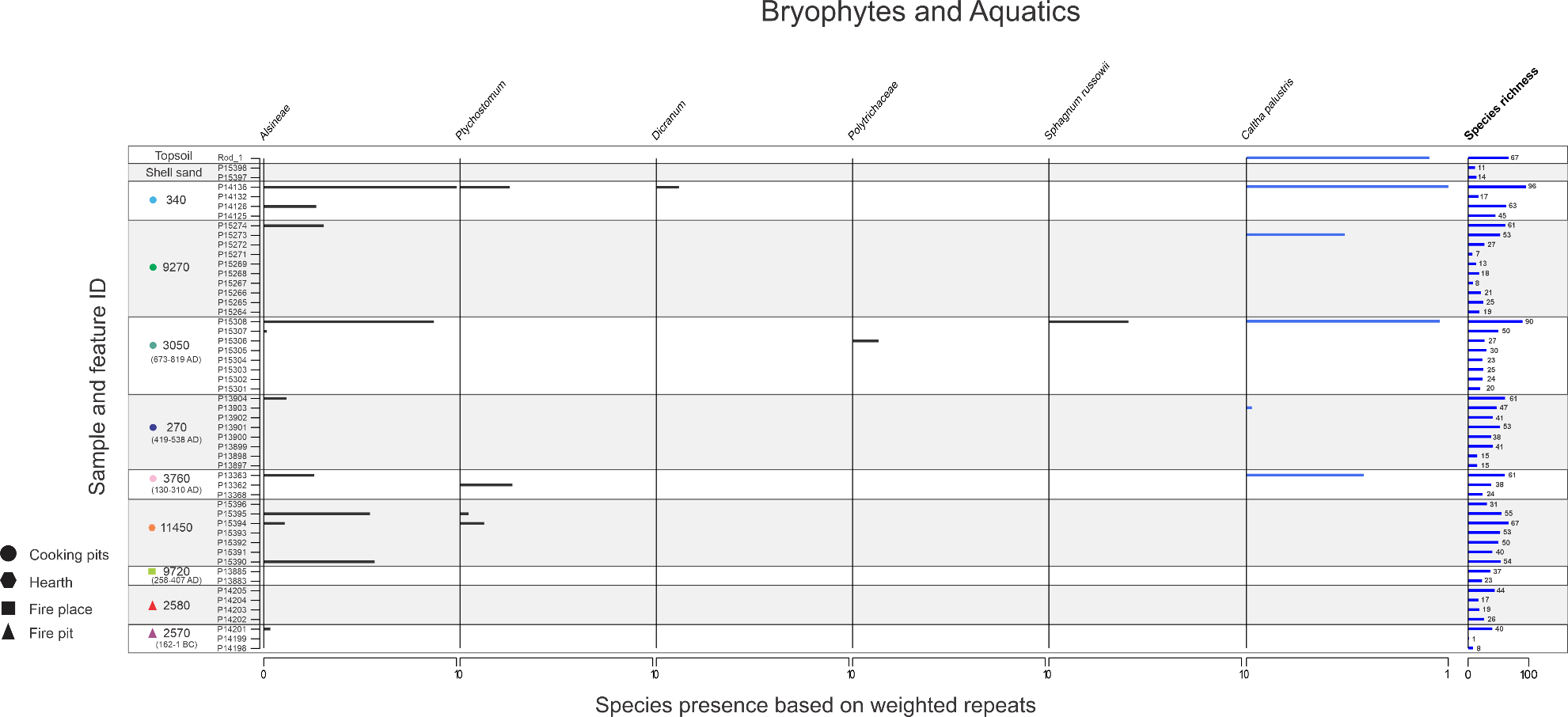

**Supplementary Figure 19.** Bryophyte and aquatic species present in samples based on weighted repeats. Species present in all eight replicates of a sample equal one. Species richness for each sample represented by blue bar.

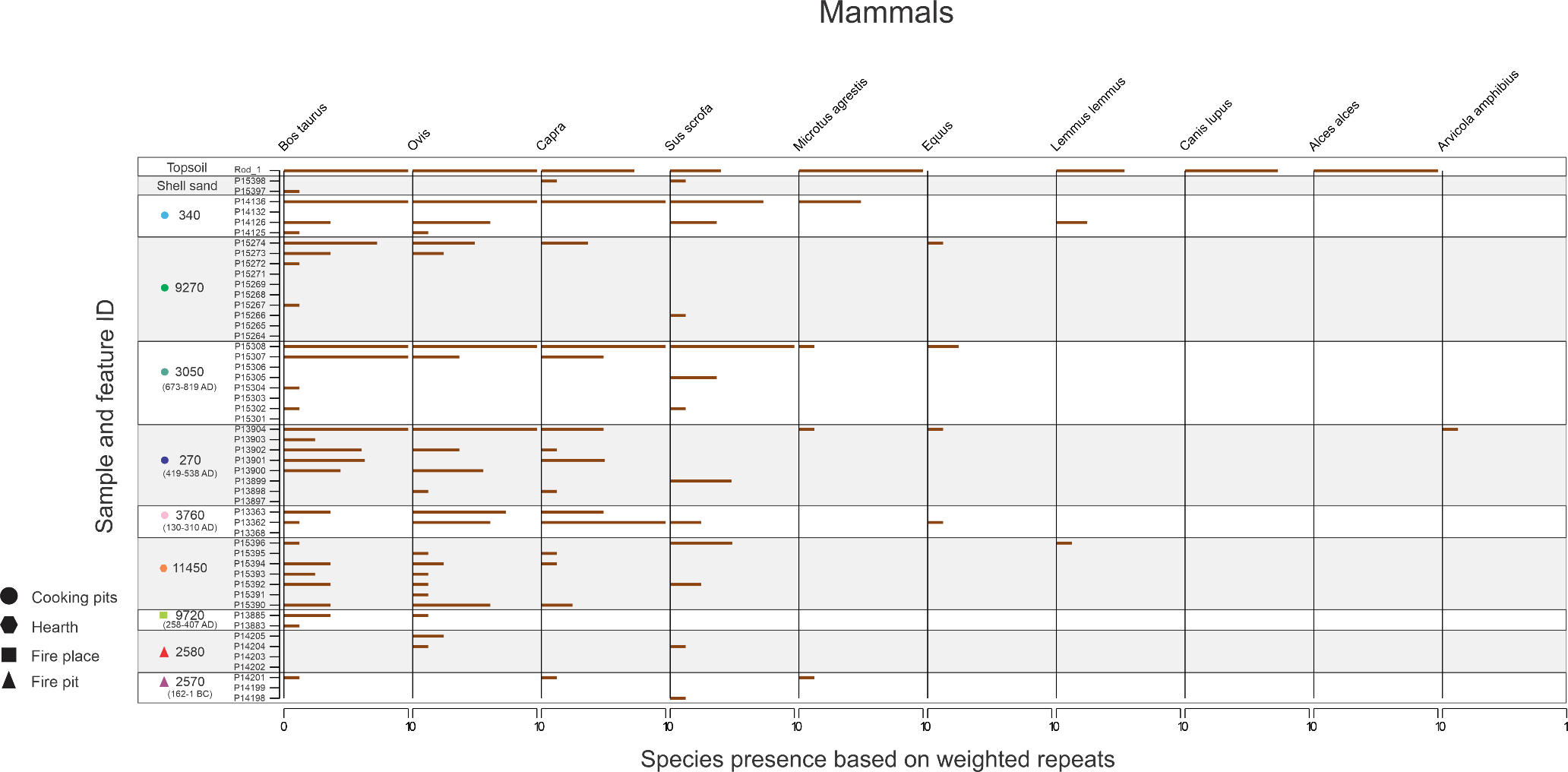

**Supplementary Figure 20.** Mammal species present in samples based on weighted repeats. Species present in all eight replicates of a sample equal one.

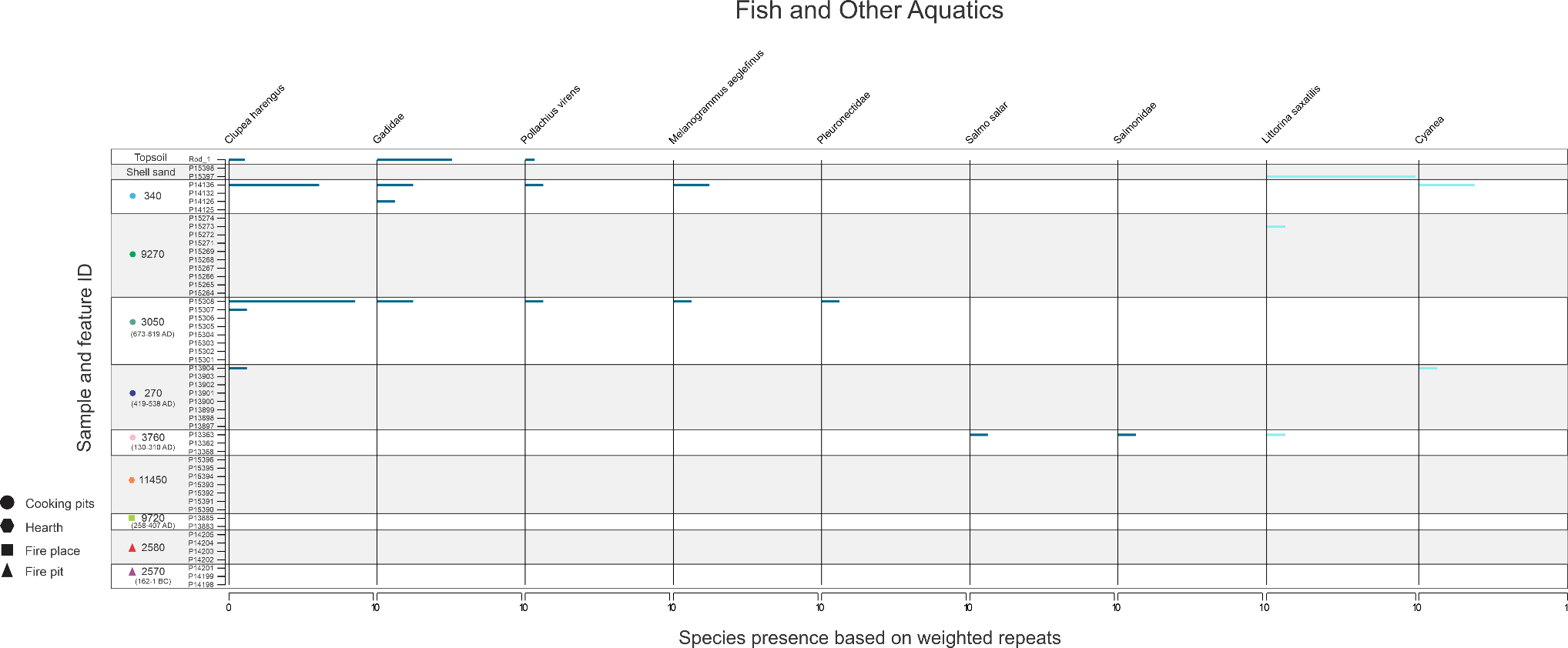

**Supplementary Figure 21.** Aquatic species present in samples based on weighted repeats. Species present in all eight replicates of a sample equal one.

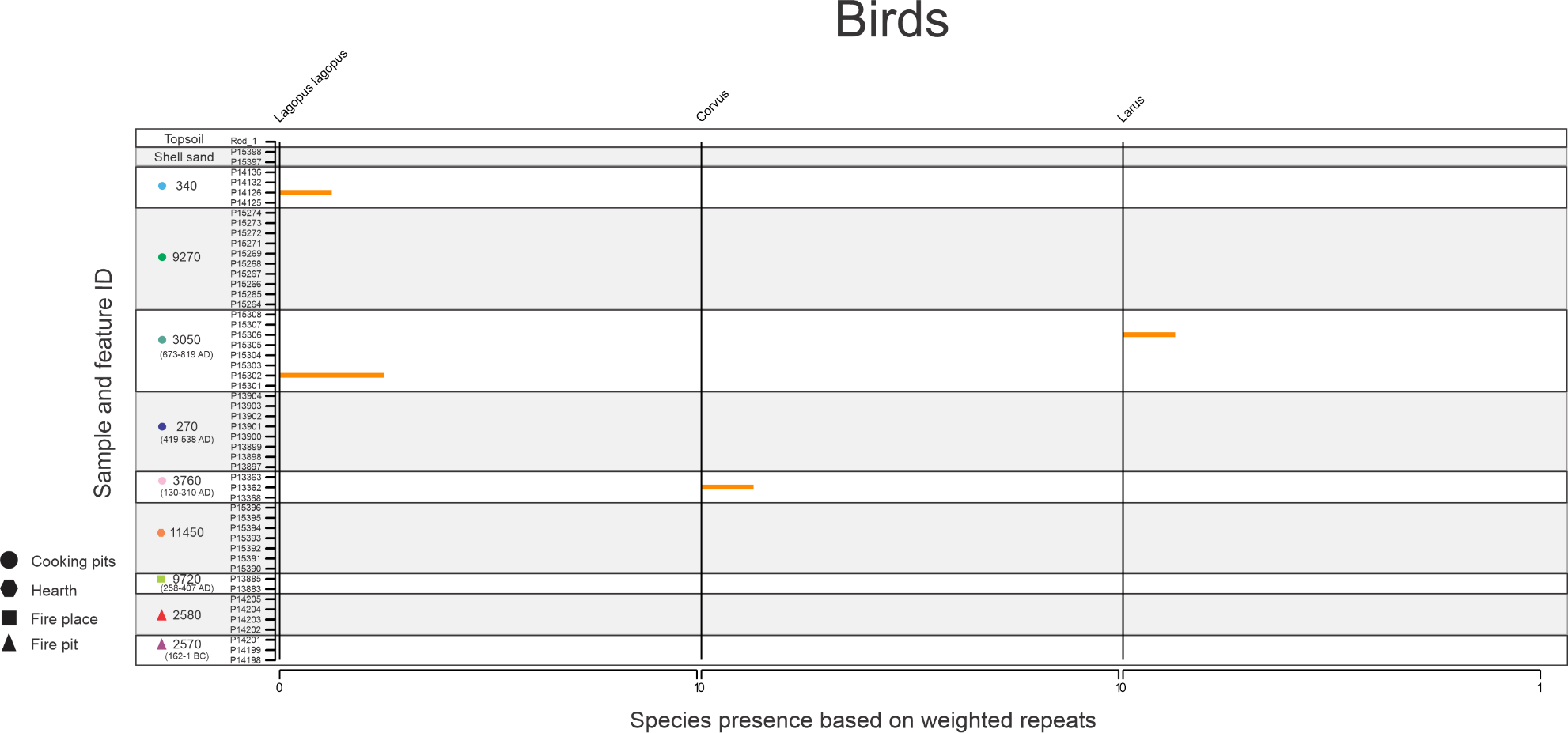

**Supplementary Figure 22.** Bird species present in samples based on weighted repeats. Species present in all eight replicates of a sample equal one.

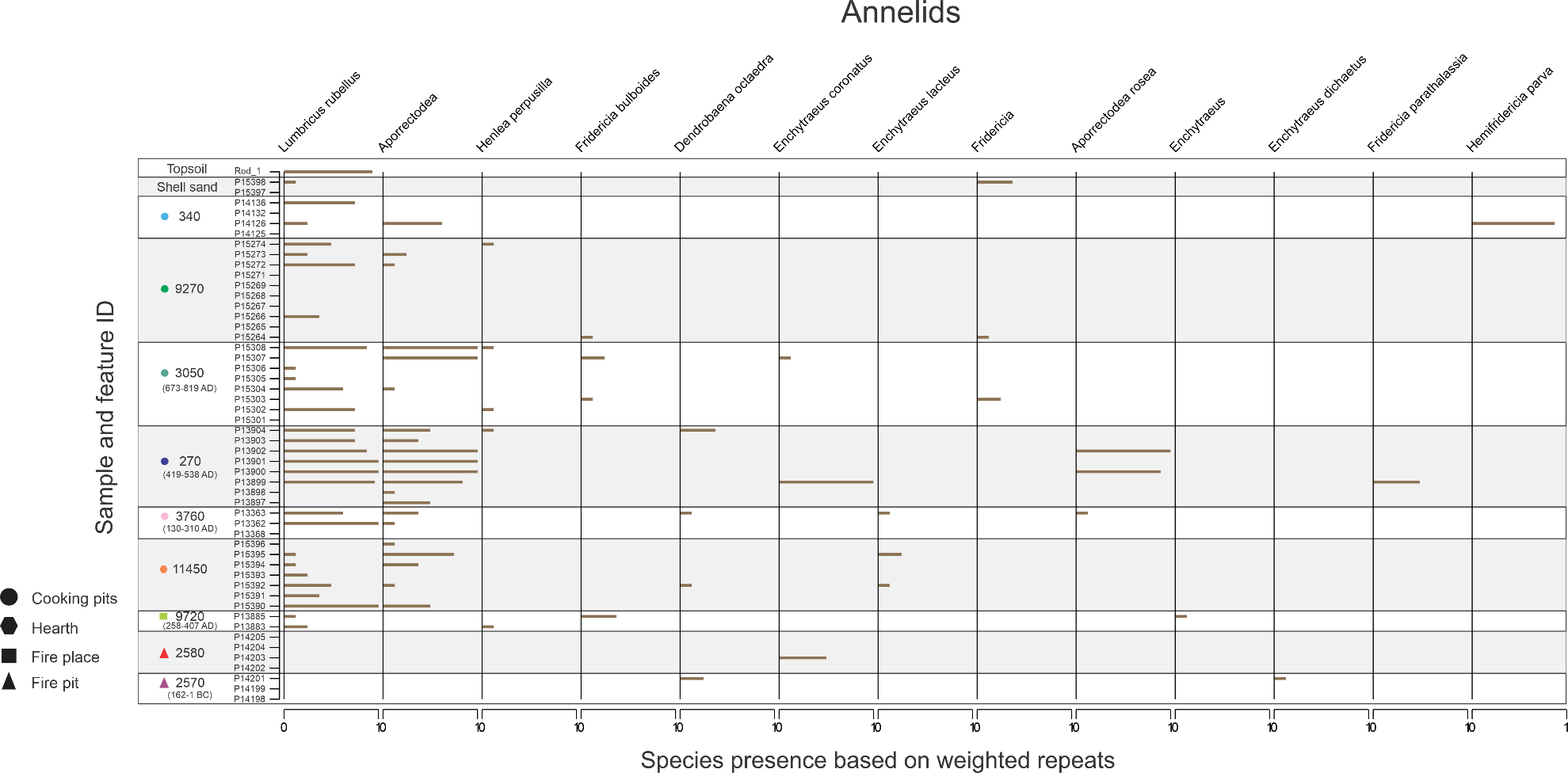

**Supplementary Figure 23.** Annelid species present in samples based on weighted repeats. Species present in all eight replicates of a sample equal one.

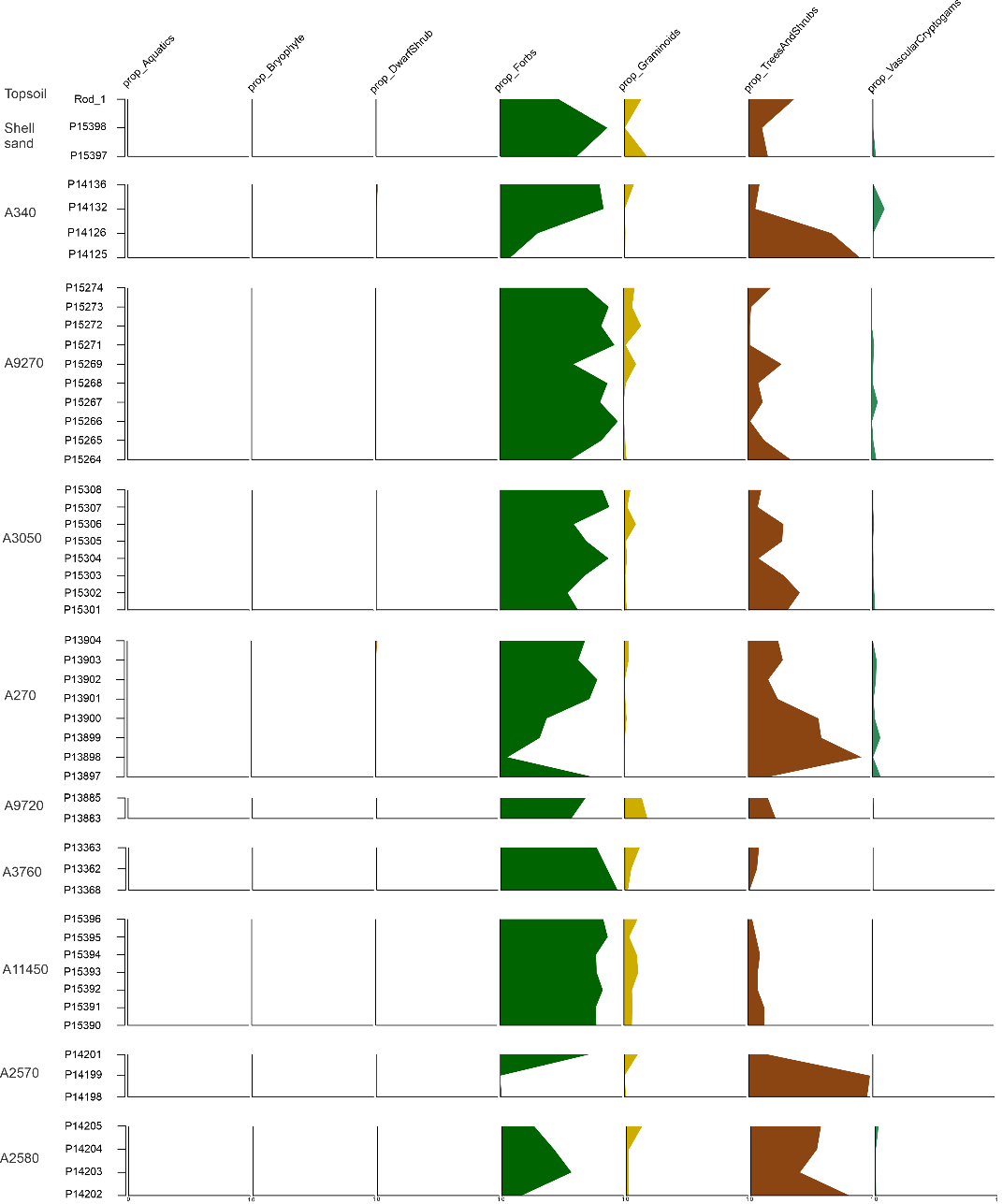

**Supplementary Figure 24.** Proportion of functional groups per sample and feature. Aquatics, bryophytes, and dwarf shrubs present but in low abundance.

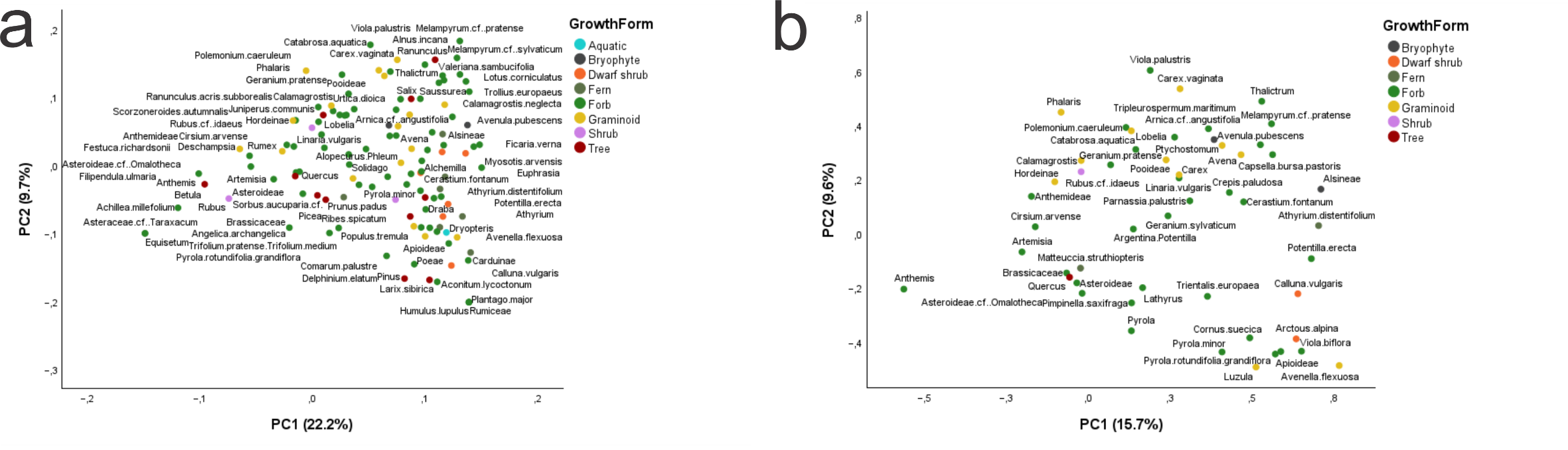

**Supplementary Figure 25.** Factor loading plots visualizing relationship of the plant taxa to the principal component analysis. **a)** plotting all plant taxa present and **b)** plotting only plant taxa found in the archaeological features but not in the topsoil.

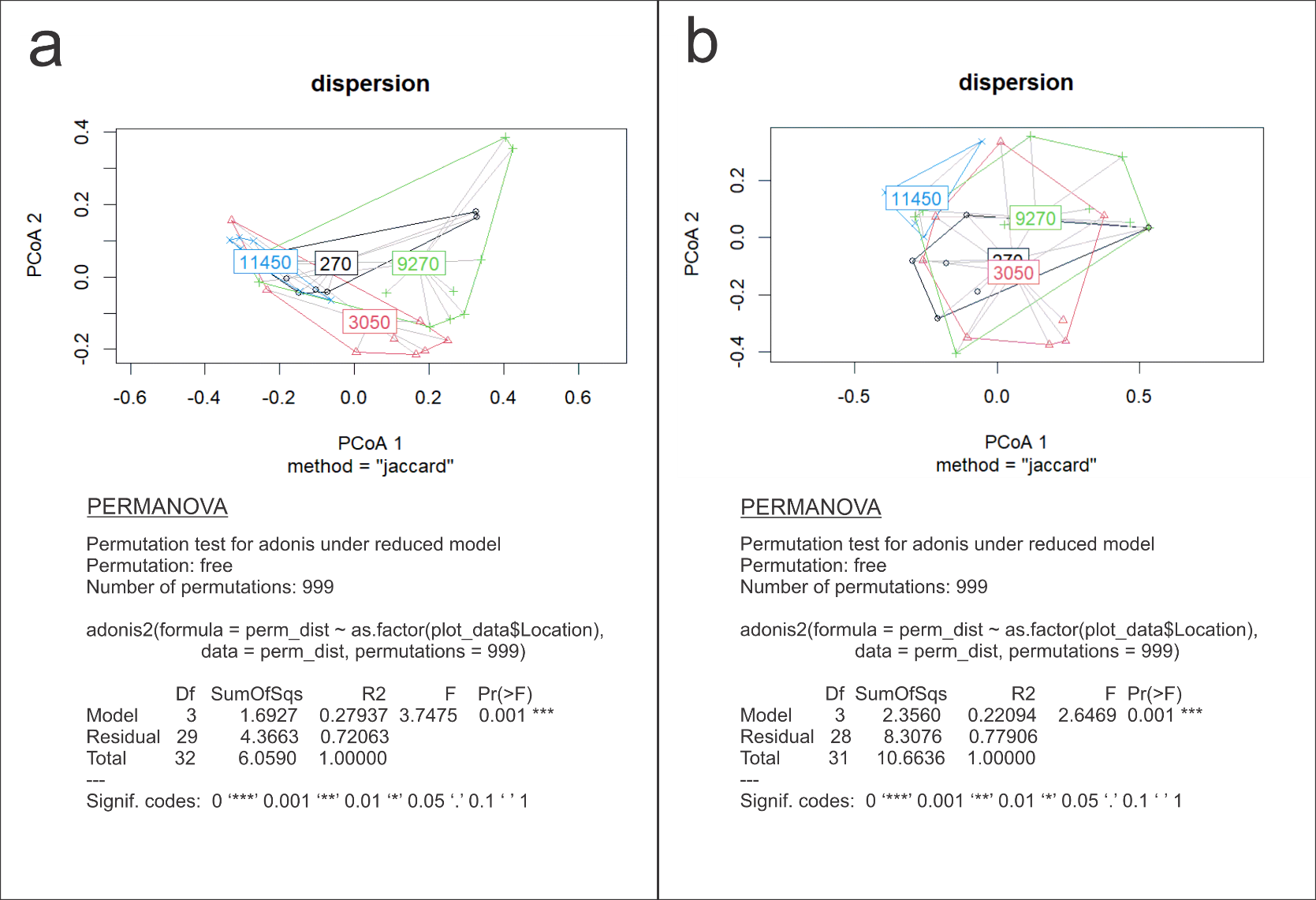

**Supplementary figure 26:** PCoA comparing cooking pit and hearth features with and without topsoil species present **a)** plotting all plant taxa present with results of PERMANOVA analysis **b)** plotting only plant taxa found in the archaeological features but not in the topsoil with results of PERMANOVA analysis.
